# Structure of SOQ1 lumenal domains identifies potential disulfide exchange for negative regulation of photoprotection, qH

**DOI:** 10.1101/2021.03.16.435614

**Authors:** Guimei Yu, Xiaowei Pan, Jingfang Hao, Lifang Shi, Yong Zhang, Jifeng Wang, Yang Xiao, Fuquan Yang, Jizhong Lou, Wenrui Chang, Alizée Malnoë, Mei Li

**Author notes:** These authors contributed equally to this work (Guimei Yu, Xiaowei Pan). Correspondence to (A.M.); (M.L.).

## Abstract

Non-photochemical quenching (NPQ) plays an important role for phototrophs in decreasing photo-oxidative damage. qH is a sustained component of NPQ and depends on the plastid lipocalin (LCNP). A thylakoid membrane-anchored protein SUPPRESSOR OF QUENCHING1 (SOQ1) prevents qH formation by inhibiting LCNP. SOQ1 suppresses qH with its lumen-located C-terminal Trx-like and NHL domains. Here we report crystal structures and biochemical characterization of SOQ1 lumenal domains. Our results show that the Trx-like and NHL domains are stably associated, with the potential redox-active motif located at their interface. Residue E859 essential for SOQ1 function is pivotal for mediating the inter-domain interaction. Moreover, the C-terminal region of SOQ1 forms an independent β-stranded domain, which possibly interacts with the Trx-like domain through disulfide exchange. Furthermore, SOQ1 is susceptible to cleavage at the loops connecting the neighboring domains both *in vitro* and *in vivo*, which could be a regulatory process for its suppression function of qH.

## Introduction

Photosynthesis is the biological process that converts solar energy into chemical energy, which sustains almost all life on earth. Oxyphototrophs often experience fluctuated illumination as light intensity and quality are constantly changing in the natural environment. Under high light or other stress conditions, illumination that exceeds photosynthetic capacity would generate reactive oxygen species, causing photo-oxidative damage to oxyphototrophs (1, 2). Thus, plants have evolved various photoprotective mechanisms, including decreasing light absorption, and dissipating excess absorbed light energy as heat (3, 4). Together, these processes are termed “non-photochemical quenching” (NPQ). NPQ is classified into four main types, namely the energy-dependent quenching, “qE”; zeaxanthin-dependent quenching, “qZ”; state transitions, “qT”; and photoinhibitory quenching, “qI” (5–8), according to their induction and relaxation kinetics and factor-dependency (5, 9). qE is the fastest process to turn on and relax, the formation of which requires a pH gradient across the thylakoid membrane (ΔpH), the photosystem II (PSII) subunit S (PsbS), zeaxanthin and light-harvesting complex (LHC) proteins (10–14). qZ and qT depend on a specific carotenoid zeaxanthin (15, 16) and an enzyme pair comprising a chloroplast kinase and phosphatase (17–19), respectively. The qI form constitutes a slowly reversible form of NPQ, which is due to the photo-oxidative damage and turnover of the D1 protein of PSII (1).

Recently, a sustained quenching component qH was uncovered (20, 21). This NPQ component occurs in light-harvesting complex II (LHCII), the peripheral antenna of PSII, and functions independently of the known NPQ factors such as PsbS, zeaxanthin, ΔpH formation or the qT-related kinase (20, 22). The plastid lipocalin (LCNP), a thylakoid lumen-localized protein, is necessary for qH to occur (20). Lipocalins mainly participate in binding and transporting various hydrophobic molecules (23). Several lipocalins from bacteria and plants participate in membrane biogenesis and repair in response to severe stress conditions by attaching to the membrane lipids (23, 24). A previous report suggests that LCNP mediates the modification of thylakoid membrane lipids, resulting in conformational changes of LHCII and quenching sites formation (20). The working model for qH is that under non-stress conditions, the SUPPRESSOR OF QUENCHING1 (SOQ1) prevents qH through inhibition of LCNP and that under stress conditions, this inhibition is alleviated leading to NPQ (20, 21). The RELAXATION OF QH1 (ROQH1) is an atypical hydrogenase-reductase located on the stromal side which is required for turning off qH (25).

SOQ1 is a 108 kDa chloroplast-localized membrane protein that contains three domains: a stromal-located haloacid dehalogenase-like hydrolase (HAD) domain followed by a transmembrane helix (TM), a lumenal-located thioredoxin-like (Trx-like) domain and a β-propeller NHL domain (21, 22). In addition, it also has a C-terminal fragment containing 159 residues (22), hereafter referred as CTD. A truncated form consisting of the TM and lumenal domains of SOQ1 is sufficient for preventing qH, indicating that the HAD domain is not involved in NPQ suppression (22). The Trx-like domain of SOQ1 belongs to the thioredoxin-like protein (TlpA)-like family (22). Thioredoxins are evolutionarily conserved ubiquitous small thiol oxidoreductases that play key roles in controlling reversible disulfide-bond formation of target proteins to induce their structural and functional switch (26, 27). The Trx-like domain of SOQ1 contains an atypical motif CCINC, similar to the classic WC(G/P)PC redox motif at the active site of thioredoxin proteins (22, 26). It was reported that single mutation of cysteines in the CCINC motif of Trx-like domain (C431S or C434S) disables suppression of NPQ (22), suggesting that SOQ1 prevents NPQ by redox-regulation of downstream target proteins. The NHL domain of SOQ1 is a member of the β-propeller branch, a structural motif that usually functions as substrate binding and protein–protein interactions (28). The NHL domain of SOQ1 is essential for qH suppression, through an as yet uncharacterized process, as indicated by a previous study that a glutamate-to-lysine mutation (E859K in the *soq1-2* mutant) in the NHL domain effectively prevents suppression of this NPQ component (22).

The detailed structure of SOQ1 is unknown, however the structure of its mammalian homolog, the human NHL repeat-containing protein 2 (NHLRC2) has been partially resolved (29). Mutations in NHLRC2 are associated with the fatal FINCA (fibrosis, neurodegeneration, cerebral angiomatosis) disease which for human variants limits life expectancy to only 1 to 2 years (30). The structure of the Trx-like and NHL domains of NHLRC2 have been determined but the C-terminal region could not be resolved. Solving the structure of SOQ1 lumenal domains (SOQ1-LD) would facilitate a more detailed understanding of its precise regulatory role in the NPQ process and essential function of NHLRC2 in human.

Here, we report four crystal structures of different truncations and mutants of the *Arabidopsis thaliana* SOQ1-LD containing different lumenal domain composition, namely the NHL domain (SOQ1_NHL_); the Trx-like domain containing Cys-to-Ser mutations plus the NHL domain (SOQ1_Trx(mut)-NHL_); the NHL domain and CTD (SOQ1_NHL-CTD_); and the loss-of-function mutant E859K of SOQ1_NHL-CTD_ (SOQ1_NHL(E859K)-CTD_). Our attempts to obtain the structure of full-length SOQ1 and SOQ1-LD were unsuccessful similarly to what has been reported for NHLRC2 structure (29); we were however able to solve the structure of the CTD. The CTD forms an independent domain in addition to the previously identified three functional domains of SOQ1, and shows high structural similarity with the N-terminal domain of disulfide bond protein D (n-DsbD). We further performed binding assay, cross-linking and molecular dynamics simulation to explore the structure of SOQ1-LD and its regulatory mechanism for qH. In addition, we analyzed lumen protein accumulation in wild type and mutant plants in both stress and non-stress conditions, and found that *in vivo* cleavage of SOQ1 occurs in both conditions. These findings not only set the stage for investigating a potential role of CTD in SOQ1 (and NHLRC2) function, but also provide a solid structural basis for in-depth exploration of the regulatory mechanism SOQ1 employs for NPQ suppression.

## Results

### The SOQ1_NHL_ domain forms a six-bladed β-propeller structure with residue E859 at its top surface

We used the complete SOQ1-LD for crystallization, however, only the NHL domain (residue N557-P907 (Fig. S1)) showed sufficiently clear electron density, while the Trx-like domain and CTD were untraceable due to missing density. Using SOQ1-LD as starting material, we could only obtain the SOQ1_NHL_ structure at 2.7 Å resolution (Fig. 1, Fig. S1B, Table S1). Prediction of the secondary structure of SOQ1-LD (31) indicated that the two fragments connecting either Trx-like and NHL domains (residue N542-P569, TN-loop) or NHL and CTD (residue G889-T923, NC-loop) (Fig. S2) form coils which are likely disordered. Therefore, we hypothesized that these two flexible loops were digested during the crystallization process. In agreement with this hypothesis, analysis of the molecular weight of SOQ1-LD protein used for crystallization and the dissolved crystals showed that the SOQ1-LD sample was partially degraded during crystallization into a fragment with a similar molecular weight to the NHL domain (36.5 kDa) (Fig. S3). This observation explains the fact that only the NHL domain could be traced in the crystal structure.

**Fig. 1.**
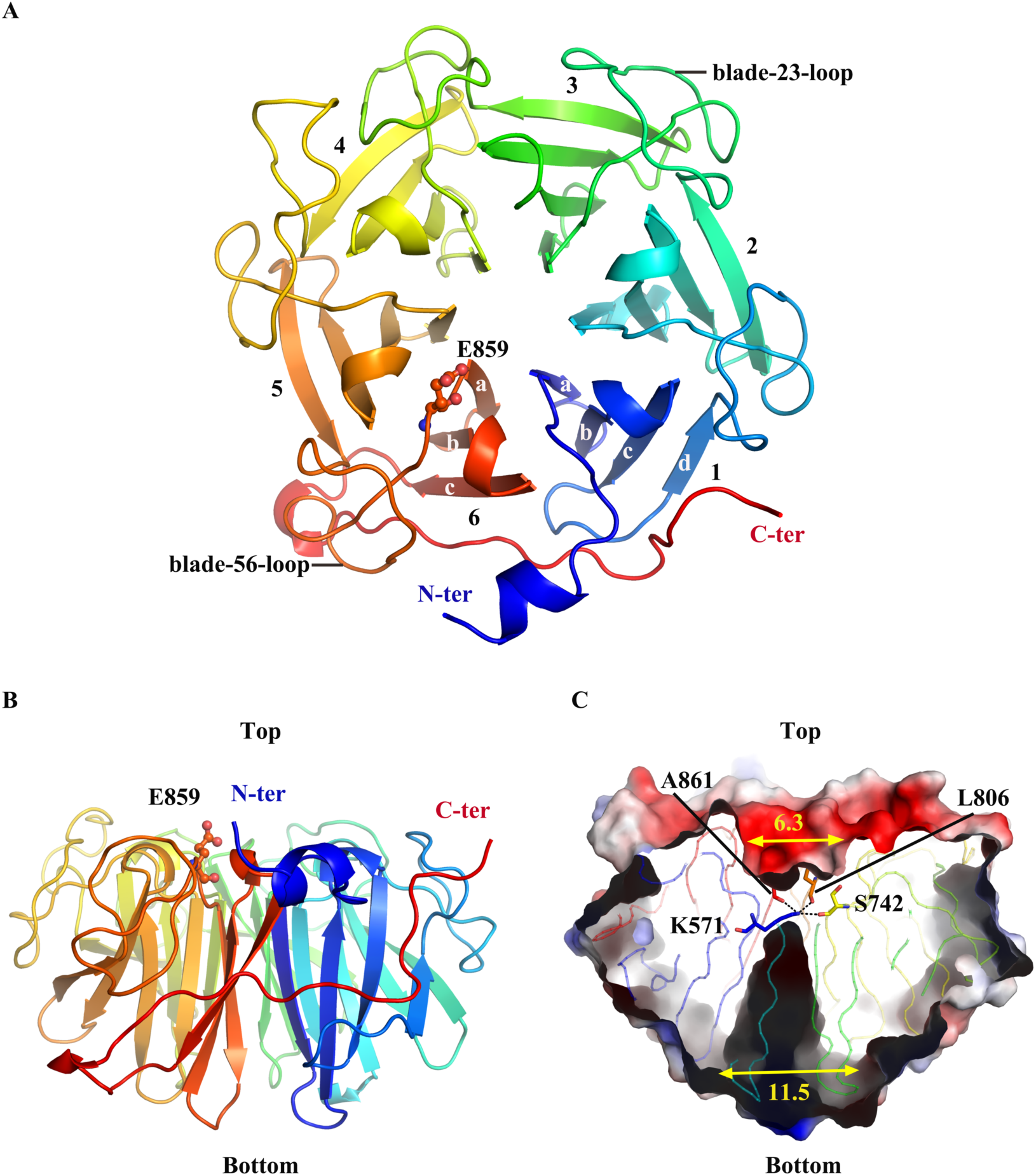
Structure of SOQ1_NHL_. (A, B) Cartoon representation of SOQ1_NHL_ in top view (A) and side view (B). The six blades of the NHL domain are numbered 1-6 from the N-terminus (blue) to the C-terminus (red). The four β-strands in one blade are labeled as a-d represented in blade 1, while blade 6 contains only three β-strands which are labeled as a-c. The residue E859 at the blade-56-loop is shown in stick-and-ball mode. (C) Side view of the discontinuous central tunnel of the NHL domain. Residues K571, S742, L806 and A861 are shown in stick mode. The hydrogen bonds are represented by black dotted lines. The diameters (Å) of two opening at the top and bottom faces of NHL domain are indicated. Surfaces are colored by their charge properties (negative in red and positive in blue).

The NHL domain of SOQ1 constitutes a six-bladed β-propeller, which we therefore named blade 1-6 (Fig. 1A). Blades 1-5 contain four classical antiparallel β-strands (strand a-d), while blade 6 is an atypical blade that consists of three antiparallel β-strands (strand a-c). This blade uses its long C-terminal tail interacting with blade 1 to enclose the entire propeller structure (Fig. 1A, B). Together, these six blades surround a discontinuous central tunnel with two openings at the surface of the NHL domain (Fig. 1C). By convention, the top face of a propeller structure is defined as the surface with a narrow opening and containing loops connecting two neighboring blades, as well as loops between the strand b and c within one blade (32). In SOQ1_NHL_ structure, the central tunnel is blocked by residue K571 from blade 1, which forms hydrogen bonds inside the tunnel with the main chain carbonyl atoms of S742, L806 and A861 from blade 4, 5 and 6, respectively (Fig.1C). As a result, the tunnel is separated into a shallow pocket opening towards the top surface and a funnel-shaped channel towards the bottom, with diameters of approximately 6.3 Å and 11.5 Å, respectively (Fig. 1C).

The six-bladed propeller located in the middle of two other domains usually serves in mediating protein-protein interactions (32). Moreover, the loops at the top surface in NHL domains generally take part in protein association and substrate binding (32). The NHL domain of SOQ1 locates between the Trx-like domain and CTD (Fig. S1). Residue E859, which is critical for proper SOQ1 function, is located at the loop connecting blade 5 and 6 (blade-56-loop) on the top surface of the SOQ1_NHL_ structure (Fig. 1A, B). This structural arrangement suggests that E859 itself associates with other domains of SOQ1 or with other proteins.

### The Trx-NHL interface harbors key residues for suppression function of SOQ1

To obtain a stable protein, we generated SOQ1-LD mutants, and used the most stable mutant form, which contains three Cys-to-Ser mutations in the redox motif of the Trx-like domain (C430S, C431S, C434S), for crystallization (Fig. S1B). The Trx-like domain was now preserved using this mutated version, but the CTD was lost during crystallization, probably due to the cleavage of the NC-loop. We thus obtained the structure of SOQ1_Trx(mut)-NHL_ (residue A393-P907) at 2.8 Å resolution (Fig. 2A, Fig. S1B, Table S1).

**Fig. 2.**
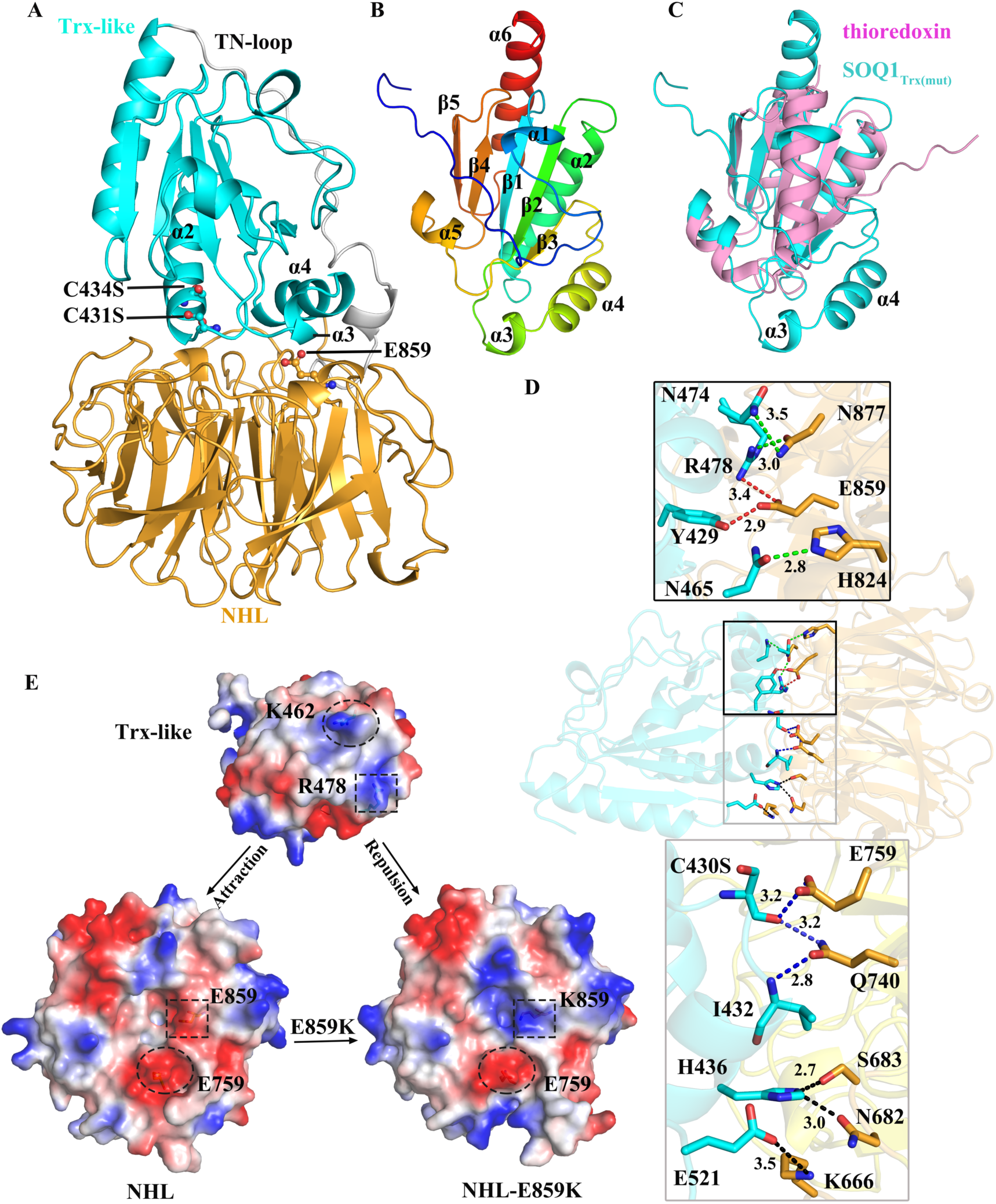
Structure of SOQ1_Trx(mut)-NHL_ and the interactions between the Trx-like and NHL domains. (A) Cartoon representation of SOQ1_Trx(mut)-NHL_ structure. The Trx-like domain and the NHL domain are colored in cyan and bright orange, respectively. The TN-loop connecting the two domains is shown in white. The C431S, C434S and E859 at the Trx-NHL interface are shown in stick-and-ball mode. (B) Cartoon representation of the Trx-like domain colored in rainbow mode. The secondary structural elements of Trx-like domain are labeled. (C) Structural comparison of Trx-like domain (cyan) and a classic thioredoxin (pink, *Arabidopsis thaliana* thioredoxin h1, PDB ID: 1XFL). Two additional α-helices in the Trx-like domain of SOQ1 are indicated. (D) Hydrogen bond interactions between the Trx-like and NHL domains. The residues involved in the inter-domain interaction are shown in sticks, with the same color codes as in (A). The hydrogen bonds are shown as dotted lines with distances (Å) labeled. The blue, green and black dotted lines represent the hydrogen bonds involving residues from the CCINC motif (C430S and I432), from the α3/α4 and from other regions of the Trx-like domain, respectively. The red dotted lines emphasize the hydrogen bonds involving E859 from the NHL domain. (E) Electrostatic potential surface of the Trx-like domain (Trx-like), NHL domain (NHL) from SOQ1_Trx(mut)-NHL_ structure and NHL domain from SOQ1_NHL(E859K)-CTD_ structure (NHL-E859K) viewed from the Trx-NHL interface. Negative potential is indicated in red and positive in blue. In SOQ1_Trx(mut)-NHL_ structure, the positively-charged residues R478 and K462 in the Trx-like domain interact with the negatively-charged residue E859 and E759 in the NHL domain, respectively. While the salt bridge between R478 and E859 is absent in the E859K mutant. The surficial regions corresponding to R478-E859/K859 and K462-E759 are highlighted by dashed boxes and circles, respectively.

The structure of the NHL domain in SOQ1_Trx(mut)-NHL_ is nearly identical to that in the SOQ1_NHL_ structure (Fig. S4). The Trx-like domain contains a central core formed by five β-strands (β1-β5) surrounded by six α-helices (α1-α6) (Fig. 2B). Unlike the classic “Trx fold’’ which is composed of five β-strands surrounded by four helices (26, 33), the Trx-like domain in SOQ1 possesses two additional short α-helices, α3 and α4 (Fig. 2C). The Trx-like domain locates at the top of the NHL central tunnel in the SOQ1_Trx(mut)-NHL_ structure. A 28-residue loop (TN-loop) links the two domains and is closely attached to both from one side (Fig. 2A). Multiple hydrogen bond interactions are formed between residues from α3, α4 and the N-terminus of α2 in the Trx-like domain and those from the top surface of NHL (Fig. 2D). Therefore, the additional α3 and α4 helices of the Trx-like domain are pivotal for the interaction between the Trx-like and NHL domains. Furthermore, the positively-charged residues R478 from α4 and K462 from α3 interact with the negatively-charged residues E859 and E759 from the NHL domain, respectively (Fig. 2E), greatly contributing to the interaction between the two domains. SOQ1_Trx(mut)-NHL_ structure confirms our suggestion that E859 in the NHL domain is involved in the inter-domain interaction. The E859K mutation (the structure is presented below) changes the local surficial charge of the NHL domain (Fig. 2E), hence affects its association with the Trx-like domain.

To explore the effect of E859K on SOQ1 conformation, we first tried to obtain the crystals of SOQ1-LD with E859K mutation (SOQ1-LD(E859K)) (Fig. S1B) but failed. Therefore we performed molecular dynamics (MD) simulation on SOQ1 Trx-NHL wild type and Trx-NHL(E859K) mutant using SOQ1_Trx(mut)-NHL_ structure as the starting model. As shown in Fig. 3, the number of hydrogen bonds between the Trx-like domain and the NHL domain of the E859K mutant form was significantly decreased, resulting in weaker interactions between these two domains. Moreover, our simulation results showed that the Trx-like and NHL domains are stably associated in the wild type protein, while separated apart in the E859K mutant (Fig. 3B, Supplementary movies 1, 2). These results confirm that residue E859 is pivotal for maintaining the compact conformation of the Trx-NHL interface.

**Fig. 3.**
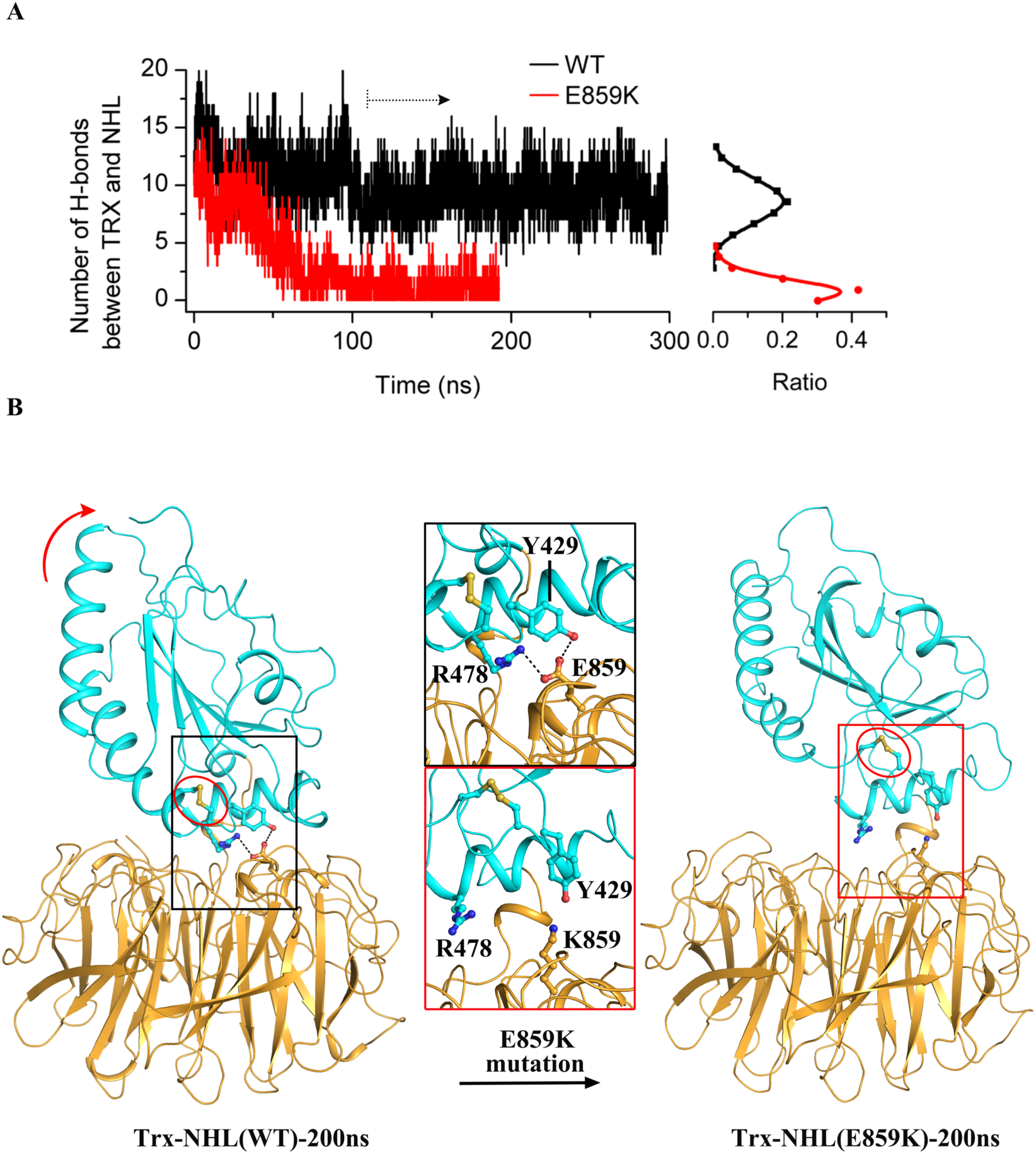
Molecular dynamics (MD) simulation shows that E859 is pivotal for maintaining the compact conformation of the Trx-NHL interface. (A) MD simulation results of the number of hydrogen bonds between Trx-like and NHL domains in the wild type (black line) and E859K mutant (red line) of SOQ1 over the simulation time duration of 300 ns and 200 ns respectively. The distribution of H-bond numbers (shown in right) was calculated by MD trajectories after 100 ns. (B) MD simulation results of the Trx-NHL domains in wild type (left) and E859K mutant (right) of SOQ1 at the 200 ns snapshot. The putative redox motif containing C431 and C434 are highlighted by red circles. Residues E859, K859, R478 and Y429 are shown as sticks. The hydrogen bonds are represented by black dotted line. The red arrow indicates the movement of the Trx-like domain away from the NHL domain in the E859K mutant of SOQ1.

The SOQ1_Trx(mut)-NHL_ structure shows considerable similarities with the recently reported structure of NHLRC2 (PDB ID:6GC1) (29). These two proteins possess the same CCINC motif in their Trx-like domain. A previous study reported that two Cys residues in this motif (C431, C434) are vital for the suppressor function of SOQ1 (22), suggesting that SOQ1 redox-regulate target proteins through this Cys pair. Although these two Cys residues were mutated to Ser in our SOQ1_Trx(mut)-NHL_ structure, they locate at the same sites and adopt conformations almost identical to those Cys residues in the reduced NHLRC2 structure (Fig. S5). The distance between C431S and C434S in SOQ1_Trx(mut)-NHL_ structure is 3.3 Å, suggesting that in the wild type SOQ1, the two Cys residues are able to form a disulfide bond through slightly conformational change under oxidized conditions in the thylakoid lumen. In addition, this motif is located at the N-terminus of α2 of the Trx-like domain, close to the Trx-NHL interface (Fig. 2A, Fig. 3B). These structural findings strongly suggest that the Trx-NHL interface comprising the CCINC motif and residue E859 is essential for SOQ1 function.

### The barrel-like CTD is likely mobile from the NHL domain

To resolve the CTD structure, we screened truncated forms of SOQ1-LD and successfully obtained a SOQ1_NHL-CTD_ protein that contains the NHL domain plus CTD, and solved its structure at 1.6 Å resolution (Fig. 4A, Fig. S1B, Table S1). The NHL domain in SOQ1_NHL-CTD_ structure adopts an identical conformation with that in the SOQ1_NHL_ and SOQ1_Trx(mut)-NHL_ structures (Fig. S4A), while the CTD comprises nine β-strands (βA-βI) connected with each other by loops, and forms a barrel-like structure (Fig. 4B). Two short continuous strands βD and βE are arranged in an antiparallel manner with βF and βC, respectively. These antiparallel β-strands further stabilize two β-sheets composed of β-strands F-B-I and C-G-H-A (Fig. 4B).

**Fig. 4.**
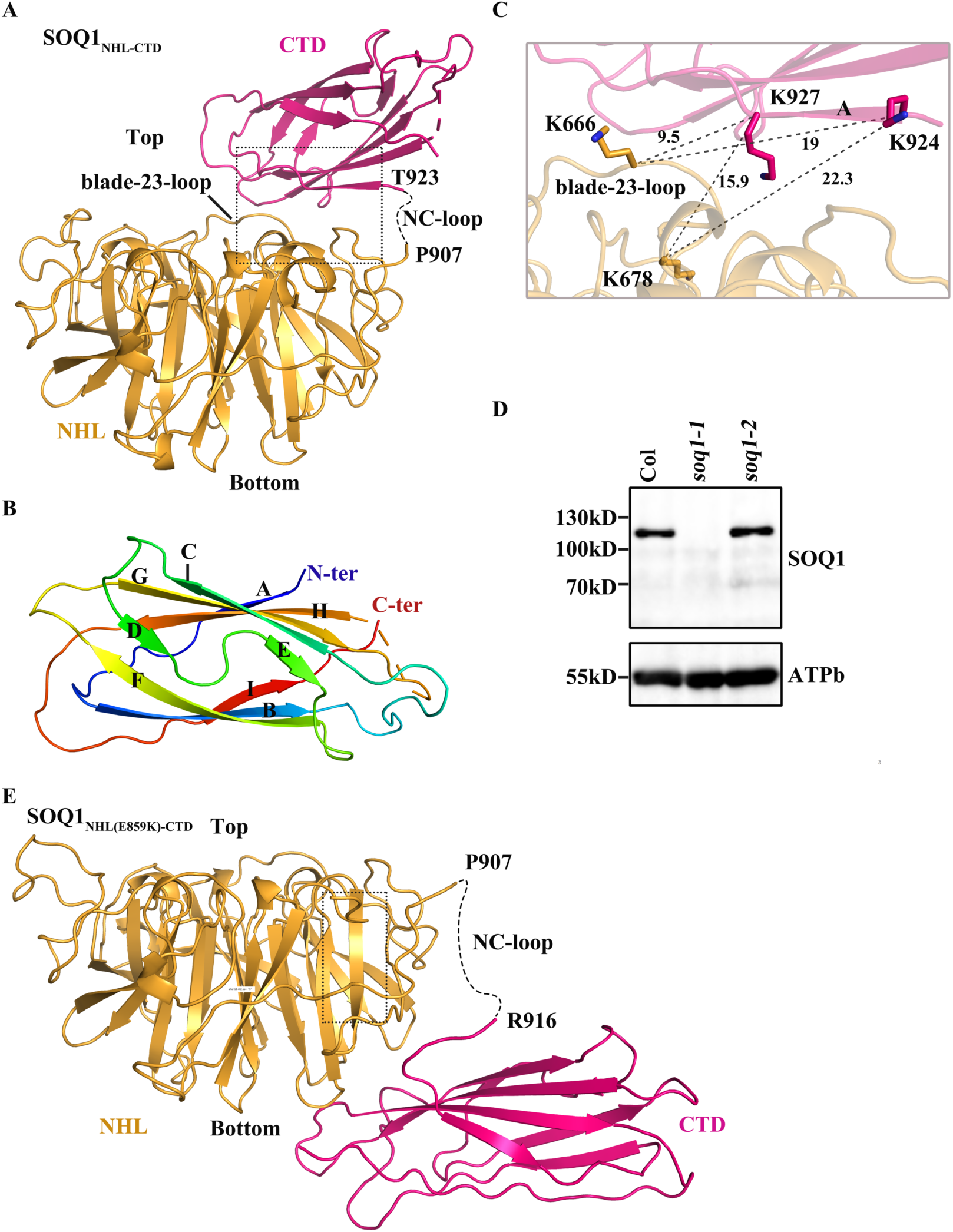
Structures of SOQ1_NHL-CTD_ and SOQ1_NHL(E859K)-CTD_. (A) Cartoon representation of SOQ1_NHL-CTD_ structure. The NHL domain and CTD are colored in bright orange and hot pink, respectively. The missing parts of the NC-loop (K908-D922) and the βG-H loop (E1008-Y1014) are shown in black and hot pink dashed lines, respectively. (B) The overall structure of CTD colored in rainbow mode. The N-terminus (blue), C-terminus (red) and the nine β-strands (A-I) of CTD are labeled. (C) The position of the cross-linked residues K666, K678, K924 and K927 (shown as sticks) in SOQ1_NHL-CTD_ structure. Each cross-linked Lys pair is connected by dotted line and the distances (Å) are labeled. (D) Total leaf proteins were separated by SDS-PAGE and analyzed by immunodetection with antibodies against SOQ1 or the β-subunit of ATP synthase (ATPb) as a loading control. Samples were loaded at the same protein content. (E) Cartoon representation of SOQ1_NHL(E859K)-CTD_ with the same color codes and in the same view (of NHL domain) as SOQ1_NHL-CTD_ structure in (A). The missing part of NC-loop (K908-L915) is shown as dashed line.

The NHL domain and the CTD of the SOQ1 structure are connected by the long NC-loop, where 15 residues were found to be untraceable due to their missing densities (Fig. 4A), probably resulting from their flexible conformation. Five symmetrically-related SOQ1_NHL-CTD_ molecules are packed closely in the SOQ1_NHL-CTD_ crystals (Fig. S6A), therefore the missing NC-loop in the structure prevents us from unambiguously determining which one of the five CTDs is indeed connected with the NHL domain. We tentatively built CTD in a position where it has the shortest distance between the last traced residue (P907) of the NHL domain and the first traced residue (T923) of CTD (Fig. S6A). However, it remains possible that other symmetrically-related CTDs are linked to the NHL domain in the SOQ1_NHL-CTD_ molecule, albeit at a longer distance.

To obtain information about the location of CTD in the SOQ1_NHL-CTD_ molecule, we cross-linked the SOQ1-LD protein with bis [sulfosuccinimidyl] suberate (BS^3^), before analyzing the cross-linked products by mass spectrometry. BS^3^ is a primary amine reactive cross-linker with a space arm of 11.4 Å that targets the amino groups on the side chains of surficial Lys residues. Considering a length of approximately 6 Å for the side chain of Lys, each Lys pair with distance shorter than 24 Å between their main chain Cα atoms can be cross-linked by BS^3^ (Fig. S6B). As shown in Table S2, K666 and K678 at the blade-23-loop of the NHL domain were cross-linked with K924 and K927 at the first β-strand of the CTD. In the built SOQ1_NHL-CTD_ structure, the CTD is attached to the top surface of the NHL domain, with its N-terminal region interacting with blade-23-loop (Fig. 4A). The distances of the K666/K678-K924/K927 pairs are ranging from 9.5 Å to 22.3 Å (Fig. 4C), consistent with our cross-linking results. In contrast, in all other symmetrically-related CTDs, K924 and K927 are located far away from either K666 or K678 of the NHL domain, with distances longer than 50 Å (Fig. S6C). These findings suggest that the built SOQ1_NHL-CTD_ structure highly resembles its native state in solution. However, two cross-linked pairs K678-K941 and K678-K952 in SOQ1-LD sample show longer distances (28.9 Å and 41.9 Å) in our SOQ1_NHL-CTD_ structure than expected for BS^3^ cross-linked pair (Fig. S7A, Table S2), suggesting that in solution, the CTD could either tilt (Fig. S7B) or be mobile to a certain degree, probably due to the NC-loop flexibility. Consistent with this suggestion, the C-terminal domain of NHLRC2 was also reported to be more disordered and flexible than the other domains of NHLRC2 molecule (29).

### The SOQ1-NHL(E859K) mutation increases mobility of CTD

A previous study showed that the SOQ1-E859K mutant (*soq1-2*) has lost its function in negatively regulating qH (22). We have first confirmed that this mutation does not destabilize the protein *in vivo* (Fig. 4D). Then we constructed the same mutant form in the SOQ1_NHL-CTD_ protein version, and solved its structure (SOQ1_NHL(E859K)-CTD_) at 3 Å resolution (Fig. 4E, Fig. S1B, Table S1). Although the NC-loop of the SOQ1_NHL(E859K)-CTD_ structure was still not completely built, we found that the densities of only eight residues were missing (Fig. S8). Therefore, we were able to identify the location of the CTD and build the SOQ1_NHL(E859K)-CTD_ structure.

The overall structures of both NHL domain and CTD in SOQ1_NHL(E859K)-CTD_ structure highly resemble those of the SOQ1_NHL-CTD_ structure (Fig. S4). However, the relative position of the two domains significantly differs between the two structures (Fig. 4A, Fig. 4E). The CTD of the SOQ1_NHL-CTD_ structure is attached on the top surface of the NHL domain, at the side of blade 2. In contrast, the CTD of the SOQ1_NHL(E859K)-CTD_ structure associates with blade 2 of the NHL domain at the bottom face, locating at the opposite side of the NHL domain compared to the CTD in the native SOQ1_NHL-CTD_ structure (Fig. 4A, Fig. 4E).

Intriguingly, neither E859 in SOQ1_NHL-CTD_ structure nor K859 in SOQ1_NHL(E859K)-CTD_ structure directly interact with CTD. In order to investigate whether the major conformational change between the NHL domain and CTD observed in the SOQ1_NHL(E859K)-CTD_ structure represents its native state in solution, we cross-linked the SOQ1-LD(E859K) protein (Fig. S1B) using BS^3^. Our results showed that most of the cross-linked pairs present in the wild type are preserved in the mutant form (Table S2), including the K666/K678-K924/K927 pairs (Fig. S9), suggesting that CTD is located at the similar position in both wild type and mutant protein in solution. However, compared with the wild type form, several additional lysine residues from the NHL domain were found to be cross-linked with K924 and K927 from the CTD in the SOQ1-LD(E859K) mutant, such as residues K828 and K859 (Table S2). Both residues are located far from the blade-23-loop of NHL domain, with the former at the bottom of NHL domain, and the latter at the blade-56-loop (Fig. S9A). These experimental findings strongly suggest that the CTD in SOQ1-LD(E859K) protein may locate in several different positions, while our SOQ1_NHL(E859K)-CTD_ structure may represent one of the several conformations, which might be more favorable for the molecular packing in the crystals. Indeed, we observed a hydrogen bond interaction between K859 and E1028 from two symmetrically–related SOQ1_NHL(E859K)-CTD_ molecules (Fig. S10), which is likely to contribute to the crystal packing. Together, our results suggest that although residue E859 is not directly involved in the interaction between the NHL domain and CTD, E859K mutation leads to the increased mobility between these two domains. Our MD simulation result showed that Trx-like and NHL domains are separated in the E859K mutant, which might affect the interaction between Trx-NHL and CTD. As a result, the three lumenal domains of SOQ1 might be weakly associated or even disordered in the E859K mutant.

### The CTD of SOQ1 is homologous to the n-DsbD

To identify the putative function of CTD, we searched for the homologous structure of CTD using the Dali server (34), and found that it adopts an immunoglobulin-like (Ig-like) fold with high structural similarity to n-DsbD from *Escherichia coli* (Fig. 5A), with a RMSD of 2.5 Å for main-chain Cα atoms. DsbD is a bacterial membrane-located protein, serving as an electron hub transferring electrons from cytoplasmic thioredoxin to periplasmic oxidoreductases (35). n-DsbD contains two essential Cys residues (C103 and C109) which can pass reducing power to substrate proteins (36). The CTD of SOQ1 also contains a corresponding pair of Cys residues (C1006 and C1012) (Fig. 5B), which are conserved among SOQ1 homologs, in the loop linking βG and βH (βG-H loop) (Fig. S2). Although residue C1012 was not built in the SOQ1_NHL-CTD_ structure (Fig. 4A, Fig. S4B), the βG-H loop is well modeled in SOQ1_NHL(E859K)-CTD_ structure, showing that the two Cys residues (especially C1012) in CTD are located at similar positions to those in n-DsbD (Fig. 5A). The two Cys residues are close to each other with a distance of 5.3 Å (Fig. S4B), and may therefore form a disulfide bond under oxidizing conditions.

**Fig. 5.**
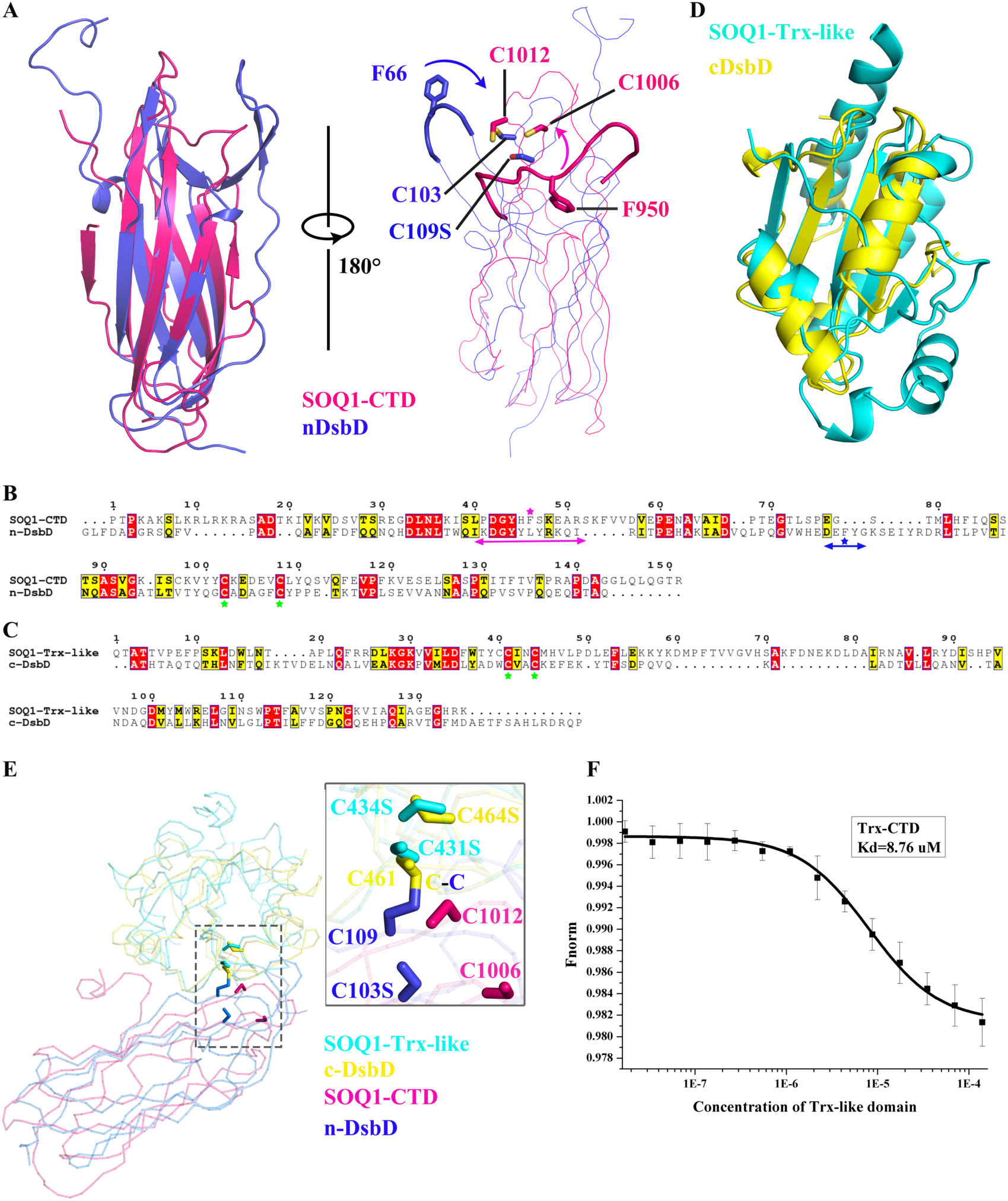
Comparison of SOQ1 with DsbD. (A) Superposition of CTD (from SOQ1_NHL(E859K)-CTD_) of SOQ1 (hot pink) with nDsbD (PDB ID: 1VRS, blue). C103, C109S and F66 of nDsbD and C1006, C1012 and F950 of CTD are shown as sticks. The blue and hot pink arrows indicate the potential movement of Cap-loop and βB-C loop for shielding the reduced C103/C109 and possibly reduced C1006/C1012 in nDsbD and CTD of SOQ1, respectively. (B, C) Sequence comparison between SOQ1 CTD and nDsbD (B), and between Trx-like domain and cDsbD (C). Conserved cysteines are labeled by green asterisks. Cap-loop of nDsbD and βB-C loop of CTD are indicated by blue and hot pink double arrows, with the phenylalanines labeled by blue and hot pink asterisks. (D) Superposition of Trx-like domain of SOQ1 (cyan) with cDsbD (PDB ID: 1VRS, yellow). (E) The CTD from SOQ1_NHL(E859K)-CTD_ structure and Trx-like domain from SOQ1_Trx(mut)-NHL_ structure are separately superposed on the nDsbD and cDsbD from the nDsbD-SS-cDsbD structure (PDB ID: 1VRS), respectively, with the same color codes as in (A) and (D). The C431S and C434S in Trx-like domain of SOQ1, C1006 and C1012 in CTD of SOQ1, C103S and C109 in nDsbD, and C461 and C464S in cDsbD are shown as sticks. (F) The binding assay of Trx-like domain with CTD measured by microscale thermophoresis (MST). MST result reveals that Trx-like domain interacts with CTD with a Kd (dissociation constant) value of 8.76 μM. Black bars represent SD. SD, Standard Deviation.

In addition, a long loop connecting βB and βC (βB-C loop) is located adjacent to the C1006-C1012 pair, and contains a phenylalanine (F950) conserved in SOQ1 (Fig. S2). This loop might correspond to the Cap-loop in n-DsbD which also contains an invariant Phe residue (F66) (Fig. 5A) and shields the Cys pair from nonspecific redox reactions (36). Thus, the CTD of SOQ1 may function similarly to n-DsbD, interacting with and passing reducing power to other substrate proteins.

### Potential redox coupling between SOQ1 Trx-like domain and CTD

A model of the complete SOQ1-LD would help to better understand the arrangement of the three lumenal domains, especially the relative positions of the Trx-like domain and the CTD, therefore we superimposed the structures of SOQ1_Trx(mut)-NHL_ and SOQ1_NHL-CTD_ aligned on their NHL domain, and obtained an entire model of the SOQ1-LD (Fig. S11). As shown in Fig. S11, the SOQ1-LD model shows a compact folding, the Trx-like domain sits right on top of the NHL domain, the CTD is above the NHL domain and beside the Trx-like domain. However, the C-terminal half of *α*2 in the Trx-like domain slightly overlap with βB and βI of CTD (circled part in Fig. S11). Moreover, K555 and K567 from the TN-loop are located at the Trx-NHL interface, and can be cross-linked with K941/K957 and K952/K957 from CTD in the SOQ1-LD sample. However, these cross-linked pairs in the SOQ1-LD model show longer distances (25.1 to 41.2 Å) than the BS^3^ linking range (Table S2, Fig. S11A), suggesting that in solution, the side of CTD containing K952 is lower and closer to the top face of NHL domain than that observed in our SOQ1_NHL-CTD_ structure, hence closer to the Trx-NHL interface. Residue K952 is located at the same side with the residues C1006 and C1012 of CTD (Fig. S11B). These findings suggest that C1006 and C1012 could move closer to the Trx-NHL interface and hence might be adjacent to the CCINC motif in solution (Fig. S11B). The possible location of CTD at the Trx-NHL interface is in agreement with our assumption that the separation of Trx-like and NHL domains in the E859K mutant leads to the highly flexible CTD.

Interestingly, our structural analysis of SOQ1 revealed that in addition to the structural similarity between CTD of SOQ1 and nDsbD, the Trx-like domain of SOQ1 shares common structural feature with C-terminal domain of DsbD (cDsbD). Hence both SOQ1 and DsbD contain two separated domains with similar fold, namely the thioredoxin-like fold (Trx-like domain of SOQ1 and c-DsbD) and Ig-like fold (CTD of SOQ1 and n-DsbD) (Fig. 5A-D), implying a possible functional resemblance. Previous reports demonstrated that n-DsbD interacts with c-DsbD, which allows n-DsbD to accept electrons from c-DsbD and further reduce substrate proteins (36, 37). The crystal structure of nDsbD-SS-cDsbD (the mixed disulfide between nDsbD and cDsbD) was reported, showing an inter-domain disulfide bond between C109 in n-DsbD and C461 in c-DsbD (37). To find out whether the Trx-like domain and CTD of SOQ1 could interact with each other in a similar way as c-DsbD and n-DsbD, we superimposed the Trx-like domain and CTD of SOQ1 with c-DsbD and n-DsbD in the nDsbD-SS-cDsbD structure (PDB: 1VRS), respectively. We found that C431S-C434S in the Trx-like domain and C1006-C1012 in the CTD of SOQ1 can be well superimposed with the corresponding Cys pair in c-DsbD and n-DsbD, respectively (Fig. 5E), implying that the disulfide exchange could occur between C431 and C1012 of SOQ1.

We further tested the binding affinity between the recombinant Trx-like domain and CTD of SOQ1 (Fig. S1C), by performing microscale thermophoresis (MST) experiment. The results showed that the two domains are able to interact with each other, with a Kd value of 8.76 μM (Fig. 5F). These structural features and biochemical assays together prove that in solution, the Trx-like domain and CTD are able to interact directly, and a redox coupling between the two Cys pairs from the two domains could occur at the Trx-NHL interface.

### SOQ1 domains can accumulate separately in the lumen

SOQ1 is easily cleaved at the TN-loop and NC-loop during crystallization (Fig. S1B, Fig. S3). Therefore, we tested in *Arabidopsis thaliana* whether SOQ1 domains could also accumulate separately *in vivo*. Previously, SOQ1 was found as a full-length protein in the thylakoid membrane enriched at the stroma margin (22). Here we further fractionated thylakoid membrane preparation in its membrane and lumen parts from both wild type (Col-0) and *soq1-1* mutant grown in standard low light conditions (LL, non-stress situation) or after exposure to a six-hour treatment at 4°C in high light (cold HL, stress situation) to induce qH. Using a SOQ1 C-terminal peptide antibody, we observed a total of six different bands in wild type sub-chloroplast fractions, which were absent in the *soq1-1* mutant and are therefore attributed to SOQ1 (Fig. 6A). The molecular sizes of these six SOQ1 protein forms were assessed as around 114 kDa, 85 kDa, 82 kDa, 65 kDa, 17 kDa and 15 kDa according to the protein marker. The bands with 114 kDa and 85 kDa molecular mass should correspond to the full-length SOQ1 (calculated molecular weight (MW) of 108 kDa) and the SOQ1 lumenal domains with TM (TM-Trx-NHL-CTD, calculated MW of 88 kDa), respectively. The presence of the 85 kDa band (faint band only found in the membrane-containing fractions) suggests that SOQ1 can be cleaved between the HAD domain and the transmembrane helix. The lower bands of 82 kDa, 65 kDa and 17 kDa were specifically observed in the lumen fraction, with molecular weights possibly corresponding to Trx-NHL-CTD (calculated MW of 73 kDa), NHL-CTD (calculated MW of 53 kDa) and CTD (calculated MW of 17 kDa), respectively. The faint lumenal band at 15 kDa should correspond to the CTD minus some amino acids at its N-terminus. We also found full length SOQ1 in the lumen fraction although there is no membrane contamination (based on lack of both membrane anchored Lhcb4 and TM-Trx-NHL-CTD in the lumen fraction, Fig. 6A). This point we cannot explain and will require more investigation.

**Fig. 6.**
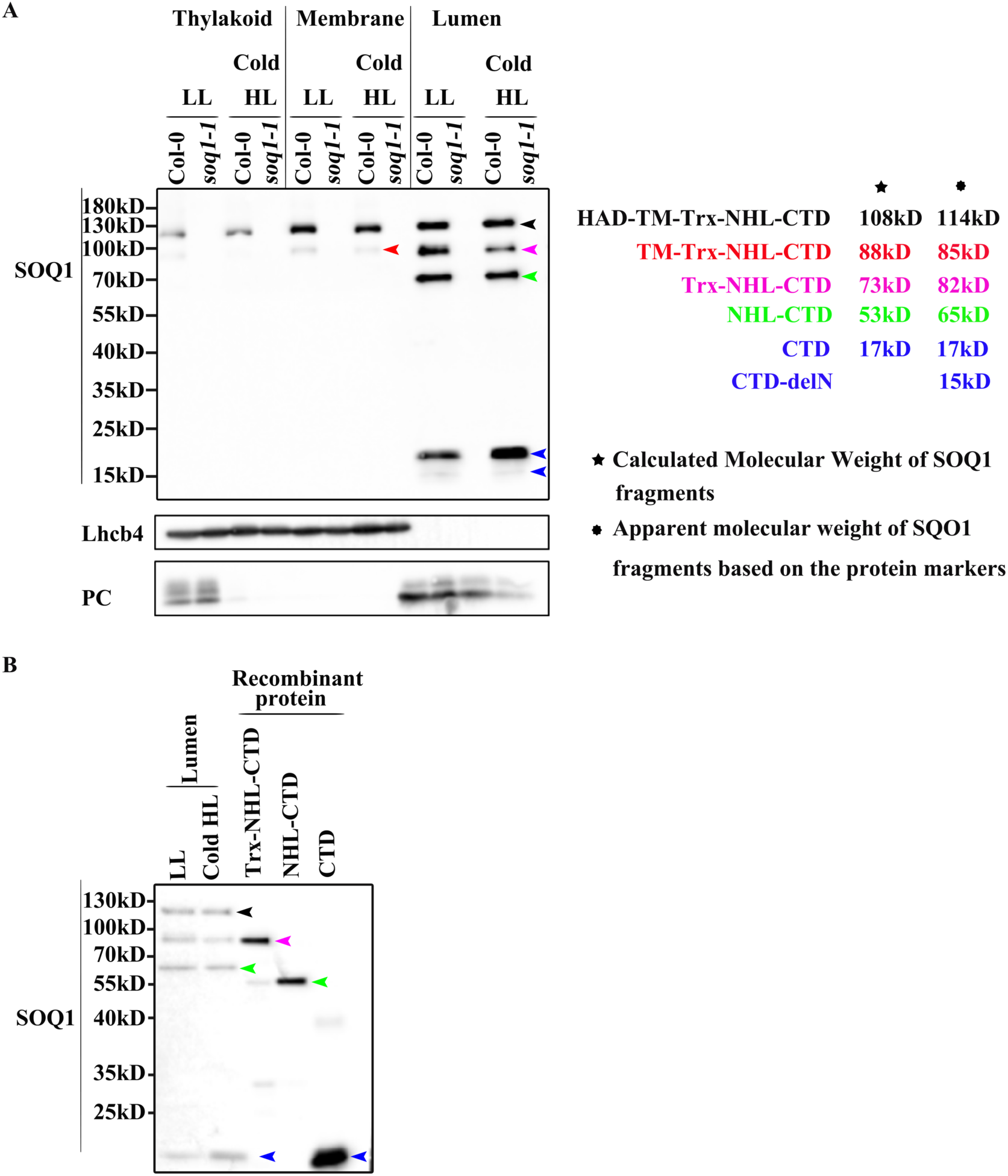
Truncated forms of SOQ1 accumulate in the thylakoid lumen. **(A)** Immunoblot analysis of chloroplast sub-fractions (Thylakoid, Membrane: thylakoid membrane after separating lumen, Lumen) of Col-0 and *soq1-1* mutant grown in standard low light condition (LL) and treated with cold and high light (Cold HL). Six bands corresponding to full-length or truncated forms of SOQ1 are detected and indicated by arrowheads with different colors. The potential fragments of SOQ1 corresponding to each band are indicated. Samples were loaded at the same protein content (3 μg). Antibodies against various proteins were used to assess the purity of the fractions. Lhcb4, thylakoid membrane light-harvesting complex b4 protein; PC, lumenal plastocyanin protein. Representative immunoblot from four independent biological experiments is shown. **(B)** Immunoblot analysis of SOQ1 lumenal truncated forms and recombinant proteins of SOQ1 with different composition of lumenal domains. Samples were loaded at the same amount of total lumenal protein (3 μg), and 0.01 pmol of recombinant protein was loaded.

In order to determine the identity of these four lower bands, we purified the recombinant proteins Trx-NHL-CTD, NHL-CTD and CTD from *E. coli* (Fig. S1D). The recombinant Trx-NHL-CTD migrated at the same size with the band at 82 kDa found in the lumen (Fig. 6B). The migration of recombinant NHL-CTD was slightly lower compared to the 65 kDa band observed in the lumen (Fig. 6B). This difference in migration sizes is likely due to the recombinant protein construct starting downstream of the *in vivo* cleavage site between the Trx-like domain and the NHL domain. The recombinant CTD (predicted size 17 kDa) migrated at the same size with the band at 17 kDa (Fig. 6B). Together, these results suggest that SOQ1 can be cleaved at the N-terminal sides of the Trx-like domain, of the NHL domain and of the CTD, and leads to the accumulation in the lumen of separate soluble SOQ1 domains consisting of Trx-NHL-CTD, NHL-CTD and CTD. The quantity of SOQ1 and these truncated forms does not seem to change between stress vs non-stress conditions (Fig. S12).

## Discussion

SOQ1 is a thylakoid membrane-anchored protein which suppresses qH with its lumenal domains (20, 22). Our structural and biochemical results showed that the three lumenal domains of SOQ1 form a compact structure, which may be critical for preventing qH. The Trx-like domain and the NHL domain are stably associated, the Trx-NHL interface harbors the key residues for SOQ1 function, and could bind the CTD. Residue E859 is pivotal in maintaining the compact conformation of SOQ1-LD. The E859K mutation changes the surface charge of NHL domain (Fig. 2E), resulting in the loose conformation of the Trx-NHL interface and hence the highly flexible CTD, thus destabilizing the three lumenal domains of SOQ1, which may lead to the loss of the suppression function of the *soq1-2* mutant reported previously (22).

In addition, our work provides new information about the CTD of SOQ1, showing that the CTD and Trx-like domain of SOQ1 adopt similar folding with n-DsbD and c-DsbD, respectively. The working mechanism of DsbD was recently proposed (36–38). The oxidized c-DsbD accepts electrons from cytoplasmic thioredoxin through the central transmembrane domain (t-DsbD), and further reduces n-DsbD through inter-domain disulfide exchange (36, 37). Afterwards, the reduced n-DsbD moves away to interact with its substrate proteins and provides the reducing equivalents (36, 37). In n-DsbD, the Cap-loop containing a conserved Phe residue is positioned adjacent to the Cys pair, shielding the thiol groups of reduced n-DsbD and facilitating the dissociation of n-DsbD from c-DsbD which prevents a futile redox cycle (36, 37). SOQ1 may function similarly as DsbD with its CTD participating in redox-transduction from the Trx-like domain to target protein(s) (Fig. 7). Our MST results showed that Trx-like domain and CTD interact in solution, which further proves the possibility of the redox coupling between Trx-like domain and CTD. We propose that the NHL domain, which shares no homology with t-DsbD and does not contain conserved Cys residues, may serve as a scaffold supporting the redox coupling between the Trx-like domain and CTD. In chloroplast, the homolog of t-DsbD is CCDA (plant homolog of the bacterial control of cell death protein), a thylakoid membrane-anchored protein which functions as a thiol disulfide transporter (22, 39). SOQ1 Trx-like domain, when not cleaved, is right after the transmembrane helix and would be localized close to the thylakoid membrane and may, therefore, accept reducing power from the stromal side through CCDA, and transfer electrons to lumenal side substrates through the CTD. The long βB-C loop containing F950 is close to the C1006-C1012 pair in CTD (Fig. 5A), and may function as a Cap-loop as that in n-DsbD. Moreover, CTD shows higher mobility compared with other domains of SOQ1 as shown by our structures and crosslinking results (Table S2), which may facilitate the interaction of CTD with its substrate proteins. Based on the structural and sequence similarity between SOQ1 and NHLRC2, we can extrapolate that the C-terminal extension in NHLRC2, with a conserved Cys pair, may also function as n-DsbD. Next it will be of interest to test whether redox imbalance in NHLRC2 potential substrates may cause FINCA disease. Conversely, knowledge on SOQ1 and NHLRC2 could inform research on finding antibiotics against DsbD as a target in human pathogens such as from the *Neisseria* species (36).

**Fig. 7.**
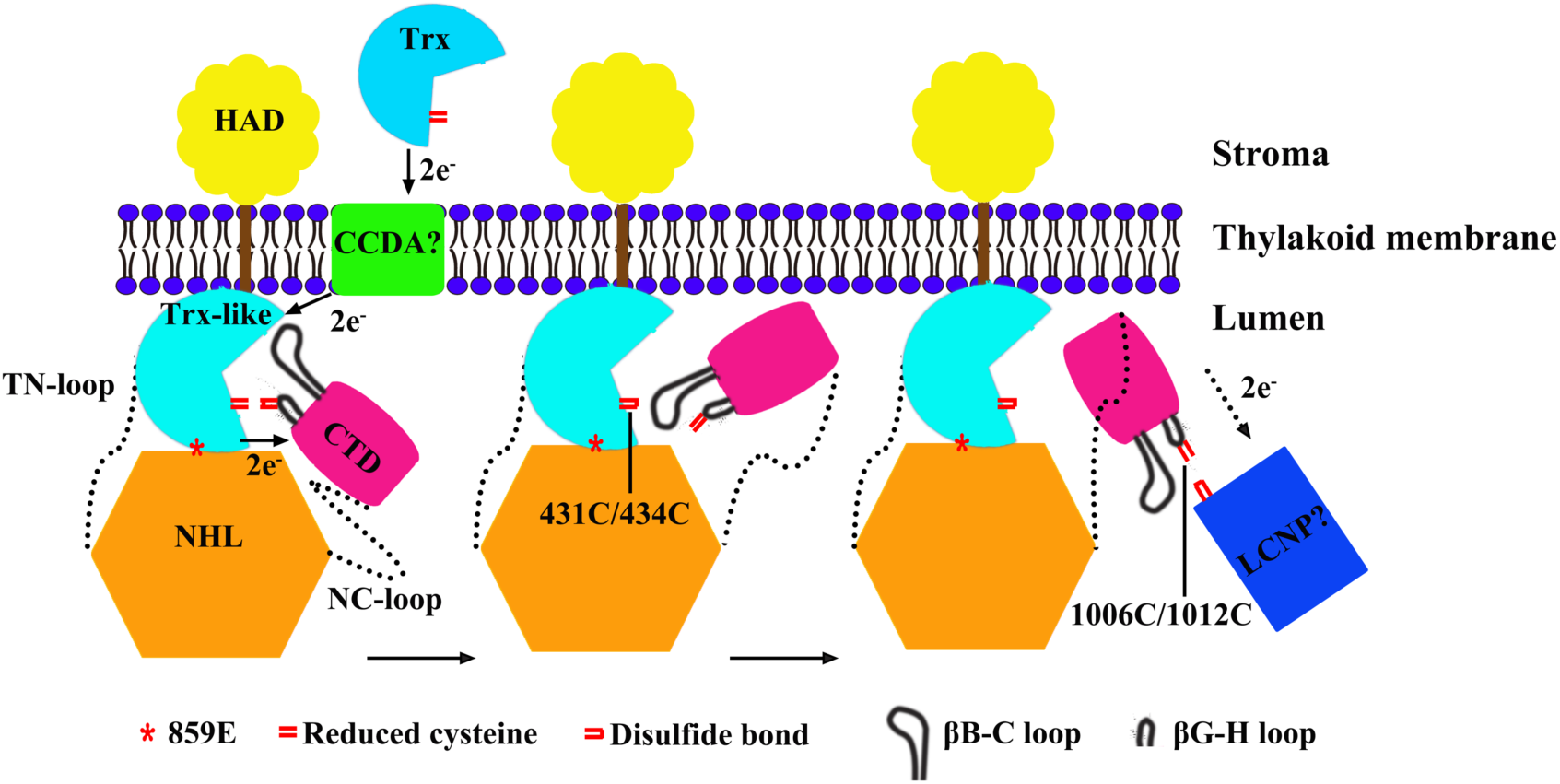
Proposed working model of SOQ1 in suppressing NPQ. SOQ1 may accept reducing power from stromal redox proteins, such as thioredoxins, through membrane-located CCDA, which changes the redox state of lumenal Trx-like domain of SOQ1, resulting in the reduced C431 and C434 pair. The CTD is located close to the Trx-NHL interface and interacts with the Trx-like domain. The Cys pairs in βG-H loop of CTD (C1006 and C1012) could be reduced by the Trx-like domain. The βB-C loop in CTD might function similarly as the cap loop in nDsbD, shielding the reduced C1006 and C1012 and inducing the dissociation of CTD from the Trx-like domain. The reduced CTD might further interact with and provide reducing power to its substrate proteins (possibly LCNP or a protein upstream of LCNP), to inhibit qH.

A potential mechanism for SOQ1 suppression of qH is through inhibition of LCNP. LCNP shows an altered electrophoretic mobility in the *soq1-1* mutant, suggesting that it might have some modifications (20). LCNP contains 12 methionines and 6 cysteines, and is located in the thylakoid lumen which is an oxidizing environment (39), thus LCNP may be activated through oxidized modifications, such as cysteine or methionine oxidation (40). SOQ1 may suppress qH by reducing the oxidized modification of LCNP directly or through other proteins which would be redox-regulated by SOQ1. Interestingly, DsbD was suggested to transfer electrons to several oxidoreductases including DsbG and PilB, to produce unmodified thiol group from sulfenylated proteins, or repair oxidized methionine residues (35). Another possibility is that SOQ1 could be involved in reduction of disulfide bonds for proper protein folding, similarly to DsbD providing electrons to protein disulfide isomerase DsbC (41). Future investigation will test whether SOQ1 functions in a similar manner as DsbD, possibly resulting in an unmodified LCNP which would be incapable of qH function.

The inhibitory effect of SOQ1 on qH may be released through SOQ1 post-translational processing, which would allow the operation of qH and protection of plants from photodamage under stress conditions. Our results showed that SOQ1 can be cleaved into distinct protein products both in chloroplast sub-fractions and *in vitro* (Fig. 6, Fig. S1B, Fig. S3). We predict the lumenal forms to be the soluble domains Trx-NHL-CTD, NHL-CTD (containing a few amino acids before the N-terminal of the NHL domain), CTD, and the 85kDa form in the membrane fractions to be SOQ1 lacking the HAD domain (Fig. 6). However, we did not observe significant differences in the accumulation of these truncated forms between non-stress and stress conditions, in which SOQ1 is hypothesized to be active and inactive, respectively (21). Whether this cleavage has a role in regulating SOQ1 function and is spontaneous or assisted by protease will require further investigation. It could be that the soluble forms of SOQ1 alter the efficiency of electron transfer from the putative membrane electron donor CCDA to the electron acceptors in the lumen. Indeed our work evidences post-translational processing in the lumen but without degradation of the sub-products which may thus have a functional role. Recently, the human SOQ1 ortholog NHLRC2 was also reported to be cleaved into a 58 kDa form (equivalent to Trx-NHL, full-length of NHLRC2 is 79 kDa) by caspase 8 in ROS-induced apoptosis (42), with a cleavage site at Asp580 before the C-terminal domain of NHLRC2 (corresponding residue is Gln903 in the NC-loop of SOQ1). It was proposed that NHLRC2 itself may participate in caspase activation by regulating the redox state of the catalytic cysteine thiol group of caspase (42). Similarly, SOQ1 could be interacting with a protease in the lumen and regulating its activity in a redox manner which would impact SOQ1 own cleavage into distinct soluble forms and possibly processing of other proteins, such as LCNP, thereby modifying their activity.

## Materials and methods

### Cloning

cDNA encoding the full-length SOQ1 and SOQ1-LD (lumenal domains) from *Arabidopsis thaliana* were synthesized (GeneScript, China) with codon optimization for yeast preferred codon. For crystallization and cross-linking, the synthesized genes and the SOQ1 truncation containing NHL and CTD (SOQ1_NHL-CTD_) were PCR-amplified and cloned into pPICZB (Invitrogen) derivative vector which contains a tobacco etch virus (TEV) protease cleavage site followed by a C-terminal green fluorescent protein (GFP) and a 6×histidine (His) tag.

During purification, we found that the SOQ1 full length protein and SOQ1-LD tend to aggregate and are potentially sensitive to protease, therefore we constructed and screened a series of cysteine-to-serine mutants of SOQ1-LD from *Arabidopsis thaliana*, including (C430S,C431S,C434S,C430S/C431S, C430S/C434S, C431S/C434S, C430S/C431S/C434S, C1001S, C1006, C1012S, C1001S/C1006S, C1006S/C1012S, C1001S/C1012S, C1001S/C1006S/C1012S). We also mutated E859 to K859 on the truncated form SOQ1_NHL-CTD_ (SOQ1_NHL(E859K)-CTD_), in order to explore the effect of E859K mutation on SOQ1. All point mutations were done through QuikChange™ Site-Directed Mutagenesis System developed by Stratagene (La Jolla, CA). The vectors were linearized with restriction endonuclease PmeI (NEB) and transformed into *Pichia pastoris* yeast GS115 strain by electroporation using Micropulser Electroporator (BioRad). Then the GS115 cell was cultured on YPDS (the yeast extract peptone dextrose with sorbitol) plate containing 400 μg/ml Zeocin at 30 °C for about 48 h.

For binding assay, the cDNA of Trx-like domain and CTD were separately PCR-amplified and cloned into pMCSG9 following the manufacturer’s instructions. Then the plasmid containing either Trx-like or CTD cDNA was transformed into *Escherichia coli* strain BL21 (DE3) (TransGen Biotech) and cultured on Luria-Bertani (LB) plate containing 100 μg/ml ampicillin.

For immunoblot analysis, the cDNAs encoding Trx-NHL-CTD (K390-R1055), NHL-CTD (P565-R1055) and CTD (E898-R1055) were amplified from pET151/D-TOPO-SOQ1 vector (41) through PCR. Then the PCR products were cloned into pET151/D-TOPO vector following the manufacturer’s instructions (the V5 epitope and TEV site were removed). The plasmids containing Trx-NHL-CTD, NHL-CTD and CTD cDNA were transformed into *Escherichia coli* strain BL21 (DE3) for protein purification.

### Protein expression in *Pichia pastoris* Yeast and purification

Ten clones of each construct were picked out from the YPDS plates, cultured in MGYH (Minimal Glycerol Media + Histidine) at 30 °C and shaken at 250 rpm. When the cell density reached OD600nm of 4.0∼6.0, the medium was changed to the MMH (Minimal Methanol + Histidine) media and the protein overexpression was induced by adding 0.5% (v:v) methanol and continually shaking at 25 °C. After about 24 h induction, 1 ml yeast cells were centrifuged at 4000 rpm and the pellet was resuspended in 200 μL distilled water. The amount of expressed proteins was monitored by the emission fluorescence value of GFP fused at the C-terminus of SOQ1 full-length or truncated proteins. The fluorescence spectra were measured at 510 nm with an excitation wavelength of 485 nm using the thermal microplate reader (Thermo). The clone with the highest fluorescence value was selected and cultured as described above. The cells were harvested and resuspended in lysis buffer A containing 25 mM Tris-HCl, pH 8.0, 0.3 M NaCl, 5% glycerol, 1 mM TCEP (tris(2-carboxyethyl) phosphine), then lysed by passing through a high-pressure crusher (JNBIO, China) at 15,000 bar for 3 times. After centrifugation at 18,000 rpm for 40 min, the supernatant was loaded onto Ni^2+^ affinity resin (Ni-NTA: GE healthcare) and washed with lysis buffer A containing 30 mM imidazole. All target proteins were eluted with 250 mM imidazole in lysis buffer A. The protein was digested with TEV protease in 8:1 (w/w) ratio at 4 °C overnight, then the mixture was loaded onto Ni-NTA affinity-chromatography again to remove the GFP and His-tag. The target proteins flowed through from the column were collected and further purified by size-exclusion chromatography (Superdex 200 10/300 GL, GE Healthcare) with lysis buffer containing 5 mM DTT (Dithiothreitol). Then the proteins were concentrated to 5∼10 mg/ml using the 30 kDa cut-off Amicon Ultra-4 centrifugal filter unit (Millipore), and stored at −80 °C for crystallization. All proteins (including SOQ1-LD, the stable mutants SOQ1-LD(M) containing C430S/C431S/C434S, SOQ1 _NHL-CTD_ and SOQ1_NHL(E859K)-CTD_) were purified with the same protocol and used for further crystallization trials.

### Protein expression in *Escherichia coli* and purification

The clones were cultured in LB medium containing ampicillin (100 μg/ml) at 37 °C and shaken at 220 rpm until the OD600 reached 0.8-1.3, then the protein expression was induced by adding 1 mM/ml isopropylthio-b-galactoside (IPTG) overnight at 18 °C. The cells were collected at 6,000 rpm for 5 min, re-suspended in lysis buffer B (25 mM Hepes, pH 7.5, 150 mM NaCl, 20 mM imidazole, 5% glycerol, 1 mM TCEP) and disrupted by sonication. The purification steps are the same with that of yeast expressed proteins but with different buffers. The wash buffer is lysis buffer B containing 10 mM, 30 mM, 50 mM, 100 mM imidazole respectively, the elution buffer is lysis buffer B containing 250 mM imidazole. The buffer for Superdex 200 10/300 GL is 25 mM Hepes, pH 7.5, 150 mM NaCl. The purified proteins were centrifuged and used for microscale thermophoresis (MST) measurement and immunoblot.

### Crystallization and structure determination

All crystals were grown using sitting-drop vapor diffusion method by mixing equal volume of protein and reservoir solution. SOQ1-LD crystals (used to solve the SOQ1_NHL_ structure) were obtained by mixing 5 mg/ml protein with a reservoir solution composed of 1.2 M NaH_2_PO_4_, 0.8 M K_2_HPO_4_, 0.2 M Li_2_SO_4_. 0.1 M Capso, pH 9.5. The crystals appeared after crystallization at 18 °C for one month. For the purpose of phasing, the heavy atom derivatives were obtained by soaking the SOQ1-LD crystals in the same reservoir solution containing 1 mM sodium salt Ethylmercuricthiosalicylic acid for 16 h at 18 °C. The SOQ1-LD(M) crystals (used to solve the SOQ1_Trx(mut)-NHL_ structure) were obtained by mixing 9.7 mg/ml protein with a reservoir solution containing 0.1 M NaCl, 0.1 M tri-Sodium citrate pH 5.6, 12% (w/v) polyethylene glycol 4000 (PEG4000), after incubating at 18 °C for about 40 days. The relatively long time for crystallization may explain the digestion of both of these protein forms.

SOQ1_NHL-CTD_ crystals were obtained at 18 °C after one week by mixing 10 mg/ml protein with the well solution containing 25-34% PEG3350, 0.2 M NH_4_Ac, 0.1 M MES, and pH 5.9-6.6. Crystals of SOQ1_NHL(E859K)-CTD_ were obtained through seeding method using grinded small SOQ1_NHL(E859K)-CTD_ crystals as seeds (43). Single crystals were grown in the condition of 25% PEG3350, 0.2 M (NH_4_)_2_SO_4_, 0.1 M MES, pH 5.3 at 4 °C after one week. All crystals were flash-cooled in liquid nitrogen before diffraction data collection.

All the diffraction data were collected at BL17U, BL18U or BL19U of the Shanghai Synchrotron Radiation Facility (SSFR)(44). The data were processed by HKL2000 program (45) and the initial phase of SOQ1-LD was obtained by the single-wavelength anomalous diffraction (SAD) method using AutoSol program in the Phenix suite (46). The partial initial model of SOQ1-LD was built by ARP/wARP and manually rebuilt in Coot (47). However, only the structure of NHL domain (SOQ1_NHL_) was solved in SOQ1-LD crystals, while the Trx-like domain and CTD might be digested during the crystallization process that lasts for one month. The structure of SOQ1_Trx(mut)-NHL_was solved by the molecular replacement method performed by Phaser in the CCP4i software suite (48), using SOQ1-LD(M) crystals. The initial models for the molecular replacement are SOQ1_NHL_ structure and Trx-like domain of NHLRC2 (PDB ID: 6GC1). Unfortunately, the CTD of SOQ1-LD(M) is also missing in the structure, only Trx-like and NHL domains (SOQ1_Trx(mut)-NHL_) are included. The SOQ1_NHL-CTD_ structure was solved by the molecular replacement method using the SOQ1_NHL_ structure as initial model, and the CTD was automatically built with Autobuild program in Phenix suite (46) and the model was adjusted manually in Coot (47). The structure of SOQ1_NHL(E859K)-CTD_ was solved by the molecular replacement method using the structure of independent NHL domain and CTD of SOQ1_NHL-CTD_ structure as initial models. All models were subjected to multiple rounds of alternative refinement with REFMAC5 in CCP4i (49) or phenix.refine in Phenix suite (46), and manual rebuilding in Coot (47).

The statistics of data analysis, phasing and structure refinement are summarized in Table S1. The quality of all the structures was checked by MolProbity (50). The molecular graphics were produced with PyMOL (http://www.pymol.org).

### Cross-linking and in-gel digestion of proteins

Because the NC-loop in SOQ1_NHL-CTD_ and SOQ1_NHL(E859K)-CTD_ structures were incompletely built due to the missing density, we cross-linked the SOQ1-LD and SOQ1-LD(E859K) proteins (1 mg/ml) with 1 mM BS^3^ ((bis[sulfosuccinimidyl] suberate) at 4 °C for 2 h, respectively, in order to determine the location of CTD. The obtained samples were analyzed by SDS-PAGE (dodecyl sulfate, sodium salt (SDS)-Polyacrylamide gel electrophoresis) and the target protein bands were cut and digested individually.

Each target protein band was cut into small plugs, washed twice in 200 μL of distilled water for 10 min each time. Then the gel bands were dehydrated in 100% acetonitrile for 10 min and dried in a speedvac for approximately 15 min. Reduction (10 mM DTT in 25 mM NH_4_HCO_3_ for 45 min at 56 °C) and alkylation (40 mM iodoacetamide in 25 mM NH_4_HCO_3_ for 45 min at room temperature in the dark) were performed, followed by washing of the gel plugs with 50% acetonitrile in 25 mM ammonium bicarbonate twice. The gel plugs were then dried using a speedvac and digested with sequence-grade modified trypsin (40 ng for each band) (Promega) in 25 mM NH_4_HCO_3_ overnight at 37 °C. The enzymatic reaction was stopped by adding formic acid to a 1% final concentration. The solution was then transferred to a sample vial for LC-MS/MS (liquid chromatography-tandem mass spectrometry) analysis.

### LC-MS/MS analysis

All nanoLC-MS/MS experiments were performed on a Q Exactive (Thermo Scientific) equipped with an Easy n-LC 1000 HPLC system (Thermo Scientific). The labeled peptides were loaded onto a 100 μm ID×2 cm fused silica trap column packed in-house with reversed phase silica (Reprosil-Pur C18 AQ, 5 μm, Dr. Maisch GmbH) and then separated on an a 75 μm ID×20 cm C18 column packed with reversed phase silica (Reprosil-Pur C18 AQ, 3 μm, Dr. Maisch GmbH). The peptides bounded on the column were eluted with a 75-min linear gradient. The solvent A consisted of 0.1% formic acid in water solution and the solvent B consisted of 0.1% formic acid in acetonitrile solution. The segmented gradient was 4–12% B, 5 min; 12– 22% B, 50 min; 22–32% B, 12 min; 32-90% B, 1 min; 90% B, 7 min at a flow rate of 300 nl/min.

The MS analysis was performed with Q Exactive mass spectrometer (Thermo Scientific). With the data-dependent acquisition mode, the MS data were acquired at a high resolution 70,000 (m/z 200) across the mass range of 300–1600 m/z. The target value was 3.00E+06 with a maximum injection time of 60 ms. The top 20 precursor ions were selected from each MS full scan with isolation width of 2 m/z for fragmentation in the HCD collision cell with normalized collision energy of 27%. Subsequently, MS/MS spectra were acquired at resolution 17,500 at m/z 200. The target value was 5.00E+04 with a maximum injection time of 80 ms. The dynamic exclusion time was 40 s. For nano electrospray ion source setting, the spray voltage was 2.0 kV; no sheath gas flow; the heated capillary temperature was 320 °C.

### Cross-linked peptides identification

The raw data from Q Exactive were analyzed with pLink2.3.7 (51, 52) for cross-linked peptides identification against target protein sequence. Some important searching parameters were set as following: trypsin was selected as enzyme and three missed cleavages were allowed for searching; the mass tolerance of precursor was set as 20 ppm and the product ions tolerance was 20 ppm; the cysteine carbamidomethylation was selected as a fixed modification, methionine oxidation as variable modifications. A 5% false-discovery rates (FDR) cutoff was set at the spectral level for peptides filter.

### Molecular Dynamics Simulations

The crystal structure of SOQ1_Trx(mut)-NHL_ (residues A391-P905) solved in this study was used as the starting model in our simulations. We manually changed S430, S431, S434 back to Cys in SOQ1_Trx(mut)-NHL_ structure to generate the Trx-NHL wild type model. The Trx-NHL wild type system was subsequently solvated in rectangular water boxes with TIP3P water model and were neutralized by 0.15 M NaCl. The E859K mutation was obtained from the same configuration using the Mutator plugin of VMD (64). The final systems contained ∼0.13 million atoms in total.

All systems were first pre-equilibrated with the following three steps: (1) 5,000 steps energy minimization with the heavy atoms of protein, followed by 2 ns equilibration simulation under 1 fs time step with these atoms constrained by 5 kcal/mol/Å^2^ spring; (2) 2 ns equilibration simulation under 1fs time step with these atoms constrained by 1 kcal/mol/Å^2^ spring; (3) 2 ns equilibration simulation under 1 fs time step with these atoms constrained by 0.2 kcal/mol/Å^2^ spring. (4) 10 ns equilibration simulation under 1 fs time step without any constrains. The resulted systems were subjected to productive simulations for more than 200 ns with 2 fs time step without any constrains.

All simulations were performed with NAMD2.13 software (53) using CHARMM36m force field with the CMAP correction (54). The simulations were performed in NPT ensemble (1 atm, 310 K) using a Langevin thermostat and Nosé-Hoover Langevin pistonmethod, respectively. 12 Å cut off with 10 to 12 Å smooth switching was used for the calculation of the van der Waals interactions. The electrostatic interactions were computed using the particle mesh Eward method under periodic boundary conditions. The system preparations and illustrations were conducted using VMD.

### Detection of the interaction between Trx-like domain and CTD by MST Assay

The Trx-like domain and CTD were separately dialyzed in a 10 kDa dialysis card (Thermo Fisher) with the Buffer (25 mM Hepes pH 7.5, 150 mM NaCl, 0.05% Tween 20) overnight at 4 °C, and then centrifuged at 135,000 rpm for 10 min. The CTD was labeled by Monolith NT.115 Protein Labeling Kit Red NHS (microscale thermophoresis grade). The Trx-like domain with a concentration of 580 μM/ml was done two-fold dilution in series with dialysis buffer, yielding the Trx-like domain samples with concentrations ranging from 580 to 8.54 nM/ml. Subsequently, 10 μL labeled CTD protein with a final concentration of 13.9 μM/ml was incubated with 10 μL Trx-like protein with different concentration and the interactions between these two domains were detected by a Monolith NT.115 instrument (Nano Temper Technologies GMBH, Munich, Germany). The MST assay was performed with 100% LED power and high MST power. The Nano Temper Analysis Software (v.1.5.41) was used to fit curves and calculate the value of the dissociation constant (Kd). Each binding assay was repeated independently four times.

### Plant material and growth conditions

Col-0, *soq1-1* and *soq1-2* mutants of *Arabidopsis thaliana* were grown in a growth chamber with 150 μmol photons m^-2^s^-1^ light intensity and 60% humidity at 20 °C during the day for 8 h and 18 °C during the night for 16 h. For cold and high light treatment, 7-week-old Col-0 and *soq1-1* mutant were illuminated in the cold room (4 °C) for 6 h at 1600 μmol photons m^-2^ s^-1^ light intensity using a custom-designed LED panel built by JBeamBio with cool white LEDs BXRA-56C1100-B-00 (Farnell).

### Lumen preparation

For lumen preparation, around 140 g leaves from 7-week-old Col-0 and *soq1-1* mutant were harvested. The method is adapted from (55). Each 20 g of leaves harvested were blended in 170 ml of homogenizing buffer (20 mM Tricine-NaOH (pH 8.4), 300 mM sorbitol, 10 mM EDTA, 10 mM KCl, 0.25% (w/v) bovine serum albumin (BSA), 4.5 mM sodium ascorbate and 5 mM L-cysteine) three times for 5 sec using a Heidolph DIAX 900 homogeniser. After filtrating through the nylon mesh, homogenate was centrifuged at 1,000 g for 2 min. Chloroplast pellet was resuspended in resuspension buffer (20 mM Hepes-NaOH (pH 7.8), 300 mM sorbitol, 10 mM EDTA, 10 mM KCl, and 5 mM MgCl_2_) using a soft brush and was centrifuged one more time for 2 min at 1,000 g to remove broken chloroplasts. Chloroplast envelopes were broken in 10 mM sodium pyrophosphate (pH 7.8) using a glass potter. Thylakoid membranes were obtained after centrifugation at 7,500 g for 5 min. Thylakoid membranes were washed once with 10 mM sodium pyrophosphate (pH 7.8), once with 300 mM sorbitol in 2 mM Tricine (pH 7.8) and once using Yeda press buffer (30 mM sodium phosphate (pH 7.8), 50 mM NaCl, 5 mM MgCl_2_ and 100 mM sucrose). Between washes, thylakoids were resuspended and homogenized by a glass potter and centrifuged at 7,500 g for 5 min. After breaking the thylakoids through the Yeda press, the thylakoid membrane fragments were ultra-centrifuged at 200,000 g for 1 h. Supernatants (lumen fraction) were moved to new tubes and concentrated at 14,000×g for 10 min at 4 °C using Amicon ultra-0.5 ml centrifugal filters (catalog number Z677094, Merck millipore, Sweden). Thylakoid, membrane and lumen protein concentrations were analyzed by Bradford assay.

### Immunoblot analysis

10 μg of total proteins from Col-0, *soq1-1* and *soq1-2* mutants or 3 μg of proteins from the chloroplast sub-fractions of Col-0 and *soq1-1* were separated on 8, 10%, or 12% SDS-PAGE followed by transfer to polyvinylidene fluoride (PVDF) membrane (Sigma-Aldrich, Stockholm, Sweden), which was then blocked with 5% (w/v) skim milk in TBST buffer (10 mM Tris-HCl, PH 7.4, 150 mM NaCl, 0.05% (v/v) Tween-20) at room temperature for 1 h. Rabbit-specific antibody against the C-terminal peptide of SOQ1 (diluted 1: 200)(20), anti-Lhcb4 (1:7,500 dilution, Agrisera, catalogue no. AS04 045), anti-RbcL (1: 5,000 dilution, Agrisera, catalogue no. AS03 037), anti-PsaA (1: 3,000 dilution, Agrisera, catalogue no. AS06 172) and anti-PC (1: 2,000 dilution, Agrisera, catalogue no. AS06 141) were added to the blocking solution, respectively, and incubation was continued 1 h at room temperature followed by three time washing with TBST buffer. The PVDF membrane was then incubated with horseradish peroxidase (HRP)-conjugated goat anti-rabbit (diluted 1:10,000; Sigma-Aldrich, Stockholm, Sweden) for 1 h at room temperature followed by three washes with TBST and detection using a Agrisera ECL kit (catalogue no. AS16 ECL-N-100). In order to identify the components of different bands, 0.01 pmol of recombinant protein was loaded as control for SDS-PAGE and immunoblot. The relative content of SOQ1 in the chloroplast sub-fractions was quantified using ImageJ based on the mean gray value of bands from immunoblots.

## Acknowledgement

We thank Matthew Brooks for providing the SOQ1 expression vector and for critical discussion together with Alexander Hertle and Krishna Niyogi. We thank Wolfgang Schröder for help with lumen preparation. We thank Yu Gao for assistance in plasmid construction and protein expression. We are grateful to the staffs at the Shanghai Synchrotron Radiation Facility (Shanghai, China) for technical support during diffraction data collection. We thank Y.Wu from Institute of Microbiology State Key Laboratory of Plant Genomics, CAS and Y.Chen from IBP for MST experiment; L. Niu, X. Ding, and M. Zhang from IBP, CAS for mass spectrometry. The project was funded by the National Key R&D Program of China (2016YFA0502900 and 2017YFA0503702), the Strategic Priority Research Program of CAS (XDB27020106), National Natural Science Foundation of China (31600609 and 31770778). X.P is sponsored by the Youth Innovation Promotion Association at the Chinese Academy of Sciences (2018128). A.M. was supported by European Commission Marie Skłodowska-Curie Actions Individual Fellowship Reintegration Panel (845687). This research (J.H and A.M.) was supported by a starting grant from the Swedish Research Council Vetenskapsrådet (2018-04150).

## Author contributions

X.P., W.C., A.M. and M.L. designed research; G.Y., X.P. and J.H. performed research with assistance from L.S. and Y.X., Y.Z. and J.L. performed the MD simulation, J.W. and F.Y. analyzed the cross-linking data. All of the authors analyzed and discussed the data, and G.Y., X.P., A.M. and M.L. wrote the paper with input from all authors.

## Supplementary Information

**Supplementary Figure S1.**
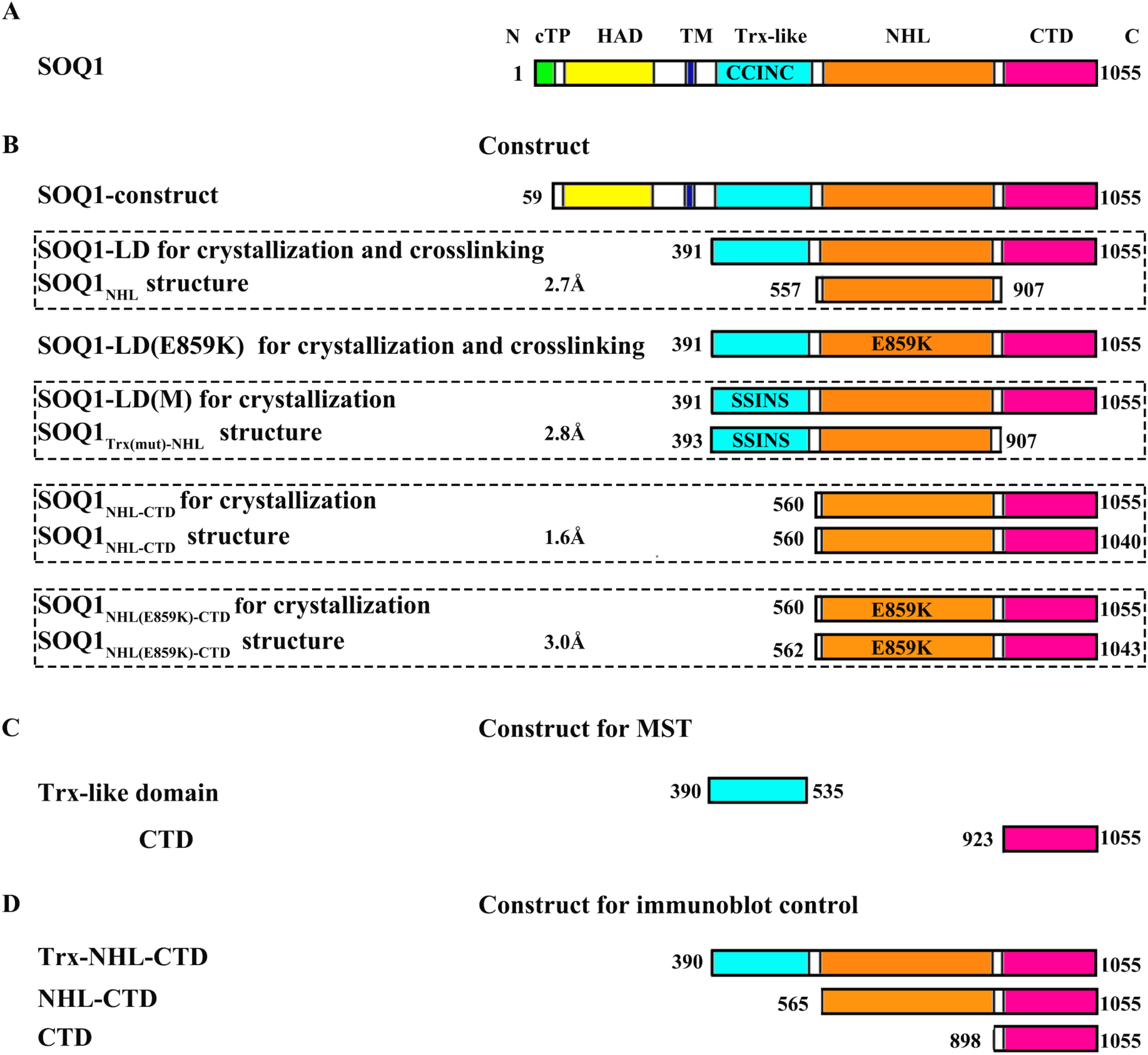
Constructs of SOQ1 used in the present study. (A) Domain structure of SOQ1. SOQ1 contains chloroplast transit peptide (cTP; green), HAD domain (yellow), transmembrane region (TM; blue), Trx-like domain (cyan) containing the CCINC motif, NHL domain (bright orange) and CTD (hot pink). Numbers indicate amino acid positions. (B) Constructs of SOQ1 truncations used for crystallization and crosslinking. The region built in each crystal structure is shown below its corresponding construct (framed in one rectangular box). Mutated residues are indicated. (C) Constructs of SOQ1 truncations for microscale thermophoresis (MST) experiment. (D) Constructs of SOQ1 truncations used as controls for heterogeneous degradation.

**Supplementary Figure S2.**
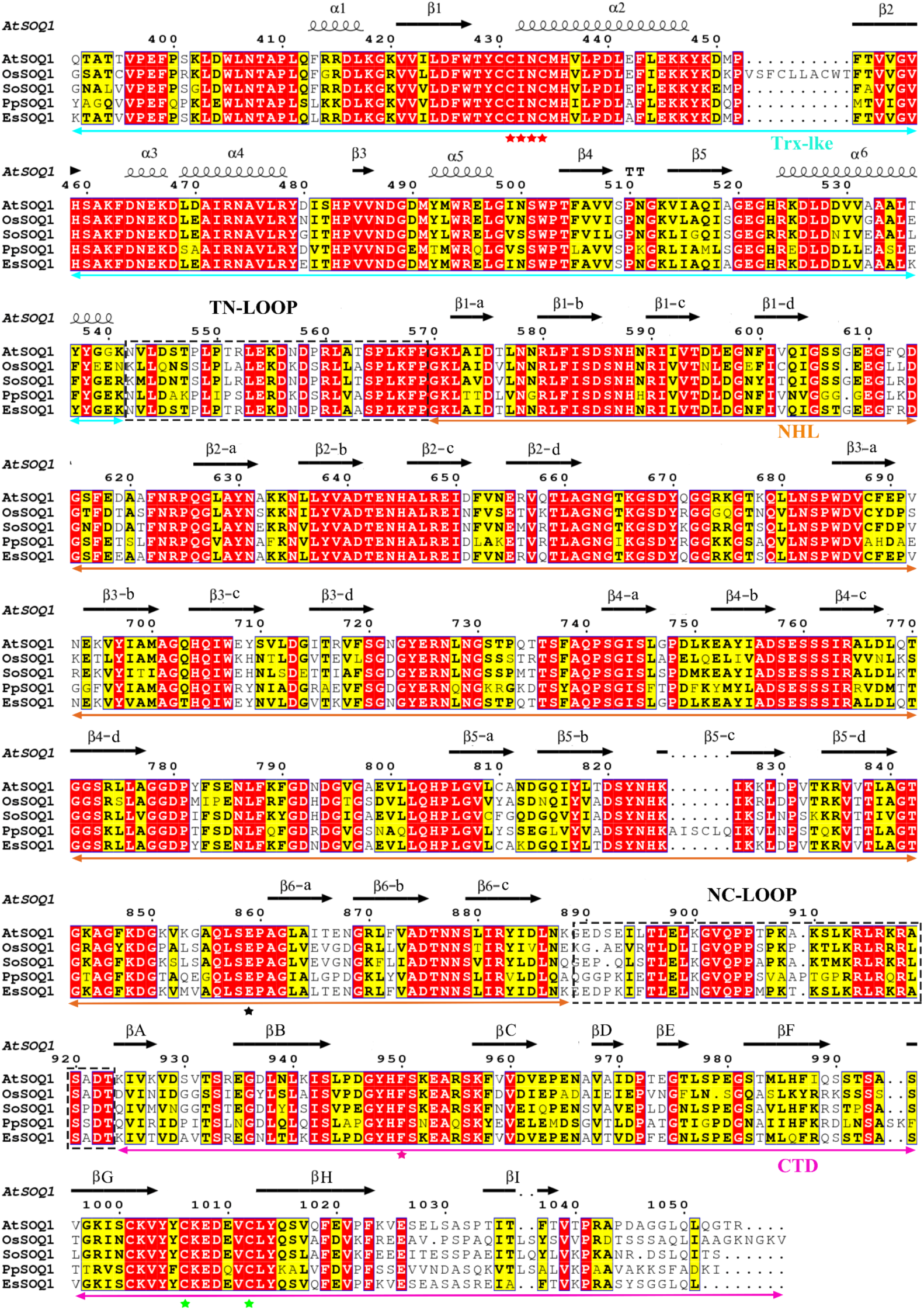
Sequence alignment of the lumenal domains of SOQ1 from various plant species. The sequences of SOQ1 lumenal part (Q391-R1055) from *Arabidopsis thaliana* |NP_564718.2|, *Oryza sativa |*XP_015631760.1|, *Spinacia oleracea |*KNA17275.1*|, Physcomitrella patens |*XP_001764907.1*|, Eutrema salsugineum |*XP_006392420.1| are labeled as AtSOQ1, OsSOQ1, SoSOQ1, PpSOQ1, EsSOQ1, respectively. The fragments corresponding to each domain and loop regions are indicated. The conserved key residues CINC in Trx-like domain, E859, F950 and C1006-C1012 pair in CTD are labeled with red, black, hot pink and green asterisks, respectively.

**Supplementary Figure S3.**
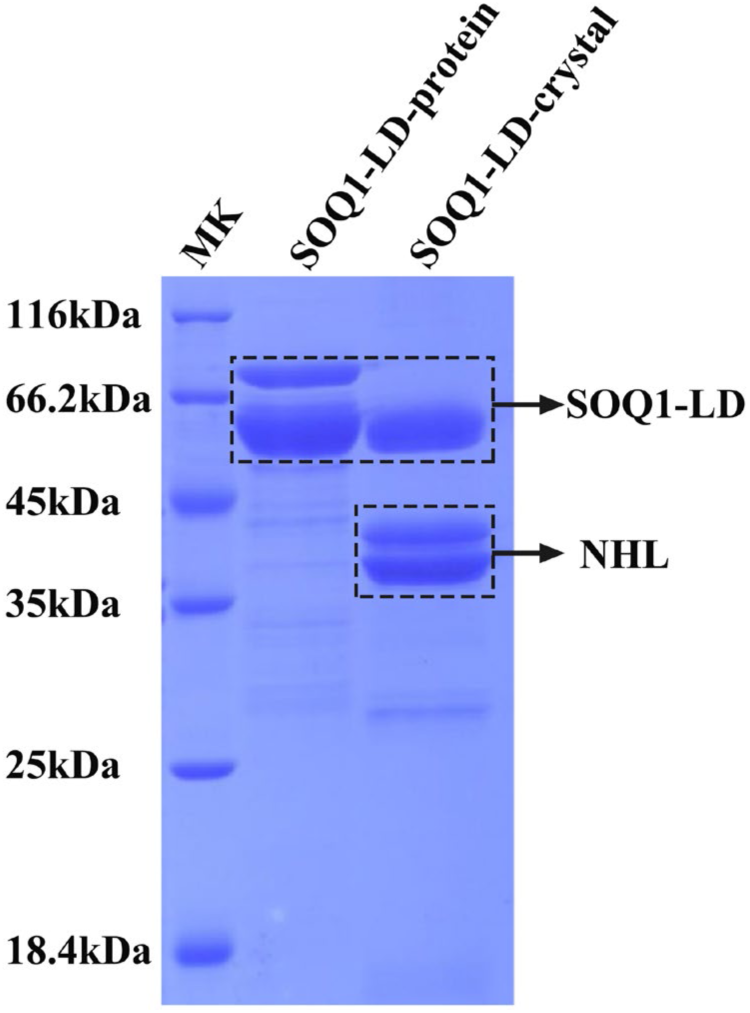
Sodium dodecylsulphate polyacrylamide gel electrophoresis (SDS-PAGE) results of SOQ1-LD protein and the yielded crystals. The protein marker (MK), SOQ1-LD protein and crystal are labeled. The bands with molecular weights similar to SOQ1-LD and NHL domain are indicated. The two bands which appear in SOQ1-LD protein are likely to correspond to the reduced (top) and oxidized (bottom) protein forms. The two bands corresponding to NHL domain may be due to heterogeneous degradation.

**Supplementary Figure S4.**
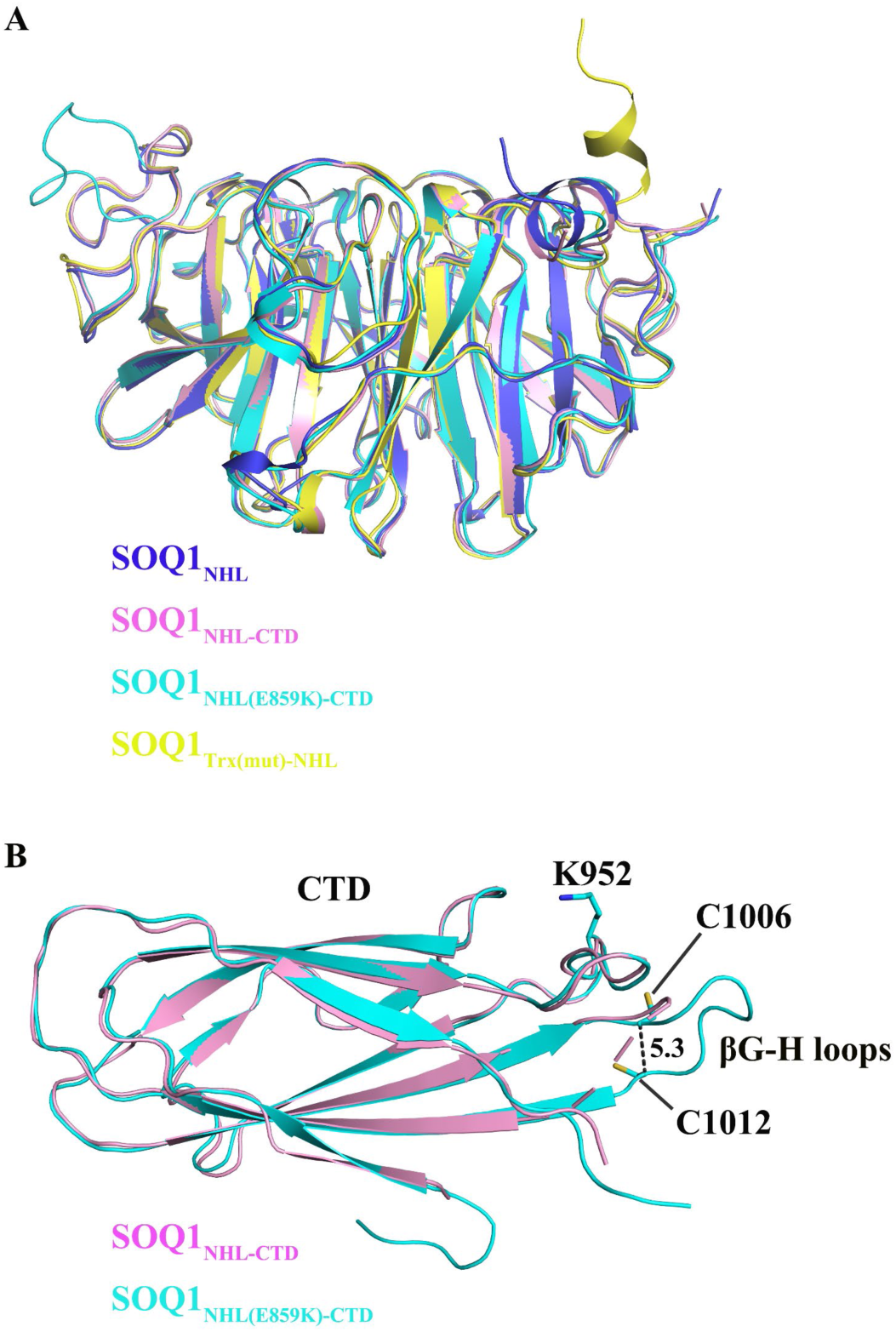
Comparison of different structures of SOQ1 truncations. (A) Structural superposition of NHL domain from SOQ1_NHL_ (marine), SOQ1_NHL-CTD_ (pink), SOQ1_NHL(E859K)-CTD_ (cyan) and SOQ1_Trx(mut)-NHL_ (yellow). The N-terminal helix as a part of TN-loop points upwards to link with the Trx-like domain in SOQ1_Trx(mut)-NHL_ structure. (B) Structural superposition of CTD from SOQ1_NHL-CTD_ (pink) and SOQ1_NHL(E859K)-CTD_ (cyan). Two Cys residues C1006 and C1012 are shown as stick mode, and the distance (Å) between them is indicated.

**Supplementary Figure S5.**
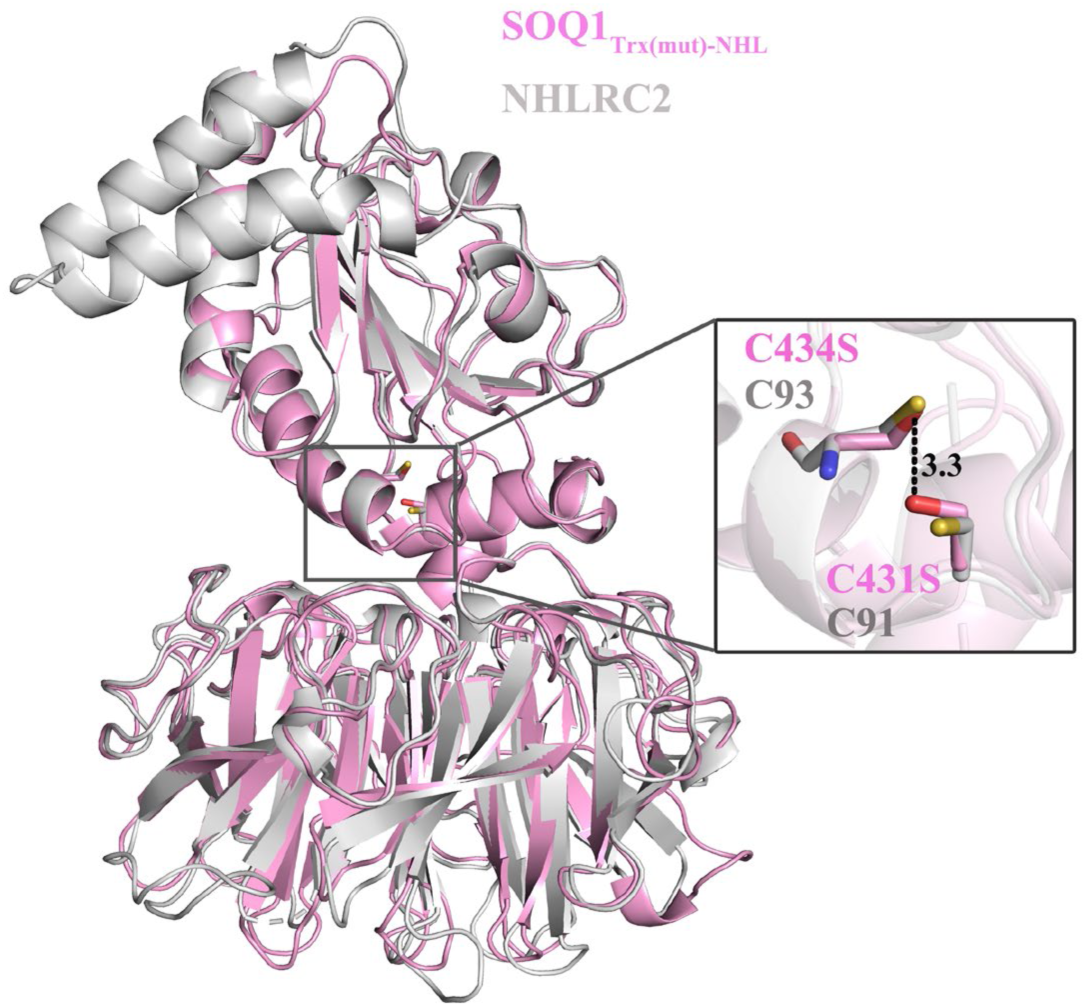
Structural comparison of SOQ1_Trx(mut)-NHL_ with reduced NHLRC2. Superposition of SOQ1_Trx(mut)-NHL_ (pink) and NHLRC2 (gray). The C431S and C434S in SOQ1_Trx(mut)-NHL_ and the corresponding Cys residues in NHLRC2 are shown as sticks. The distance (Å) between C431S and C434S is labeled. NHLRC2 shows two additional helices at the N-terminal region compared with SOQ1_Trx(mut)-NHL_. This corresponding fragment is present in SOQ1 sequence, but was not included in SOQ1_Trx(mut)-NHL_ construct.

**Supplementary Figure S6.**
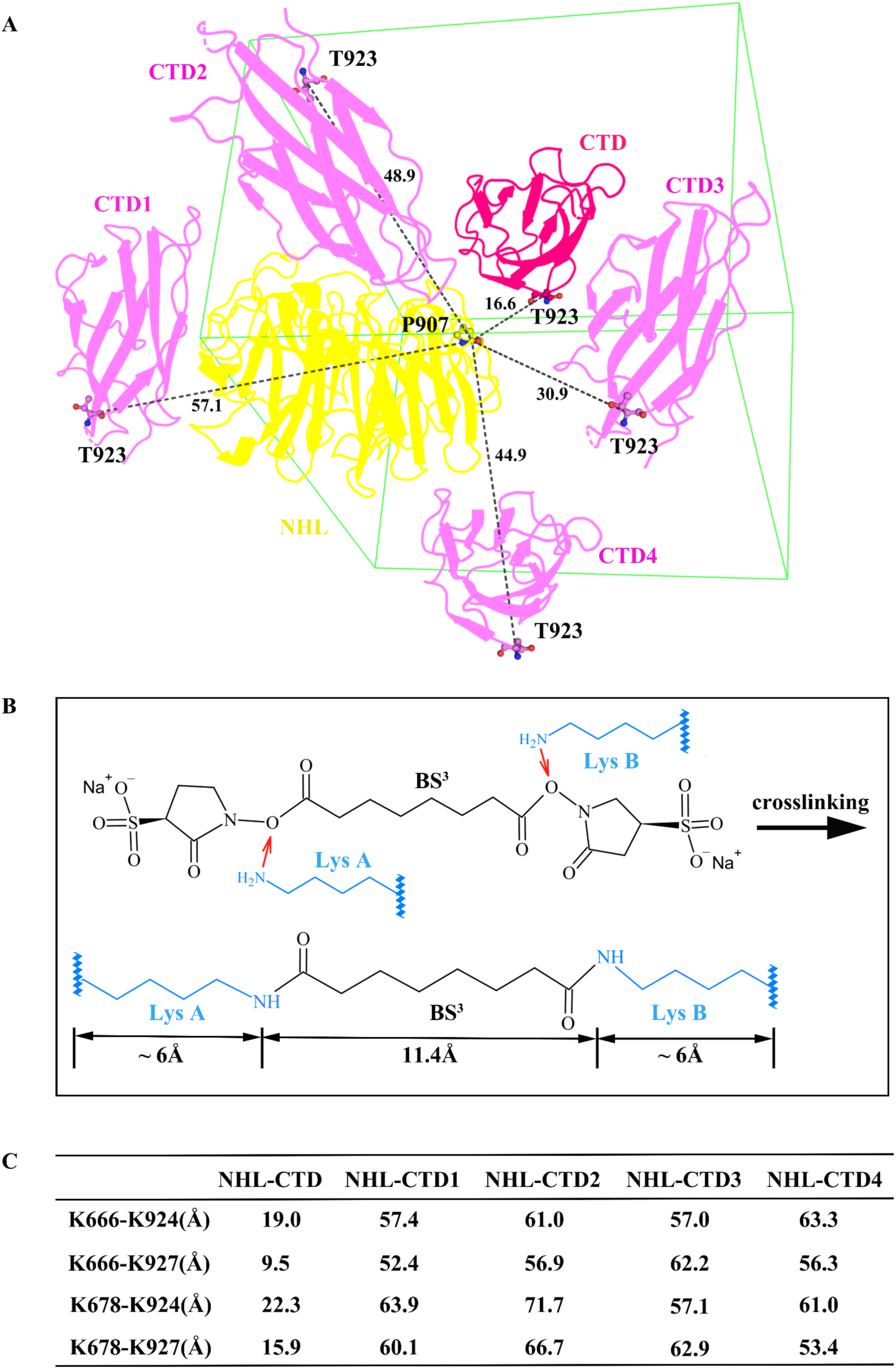
Identification of the position of CTD in the SOQ1_NHL-CTD_ structure. (A) The five symmetrically-related CTDs around one NHL domain in SOQ1_NHL-CTD_ crystal. The NHL domain is shown in yellow. The CTD built in SOQ1_NHL-CTD_ structure is shown in hot pink, and the other four symmetrically-related CTD molecules are shown in pink. The last traced residue (P907) in NHL domain and the first traced residue (T923) in CTDs are shown in stick-and-ball mode, and the distances (Å) between these two residues are indicated. (B) The scheme of BS^3^ crosslinked Lys pair. The BS^3^ is able to crosslink two Lys residues with the distance between their Cα atoms of approximately 24 Å. (C) The distances of cross-linked Lys pairs (K666/K678 from the NHL domain and K924/K927 from the five CTDs).

**Supplementary Figure S7.**
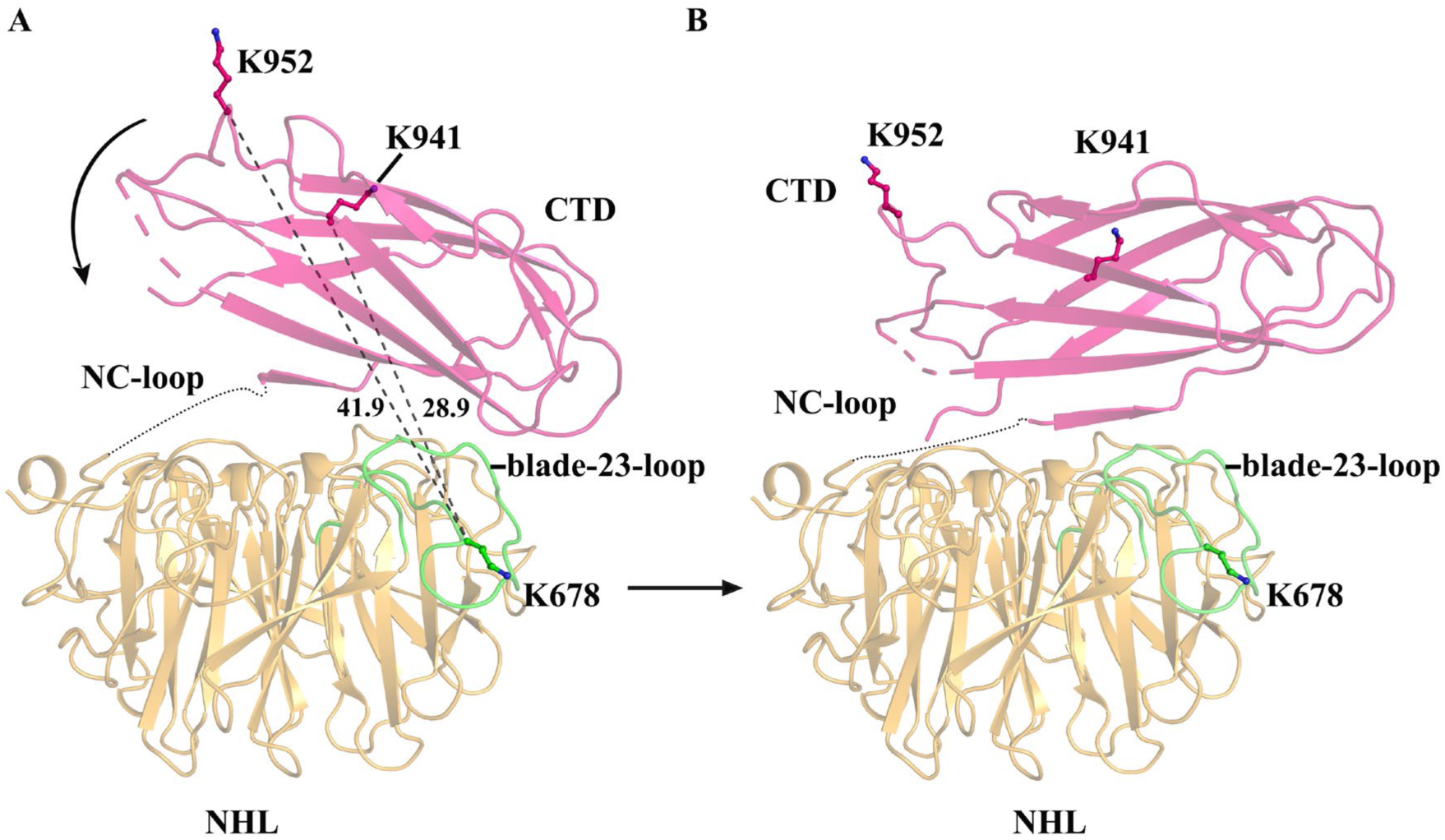
The location of cross-linked Lys residues in SOQ1_NHL-CTD_ structure. (A) The NHL domain and CTD are shown in bright orange and hot pink, respectively. The blade-23-loop of NHL domain is colored green. The cross-linked pairs with longer distances are linked by black dashed lines, and the Lys residues are shown in stick-and-ball mode. The distances (Å) between two Lys residues are indicated. (B) The modified model of SOQ1_NHL-CTD_ based on the cross-linking results.

**Supplementary Figure S8.**
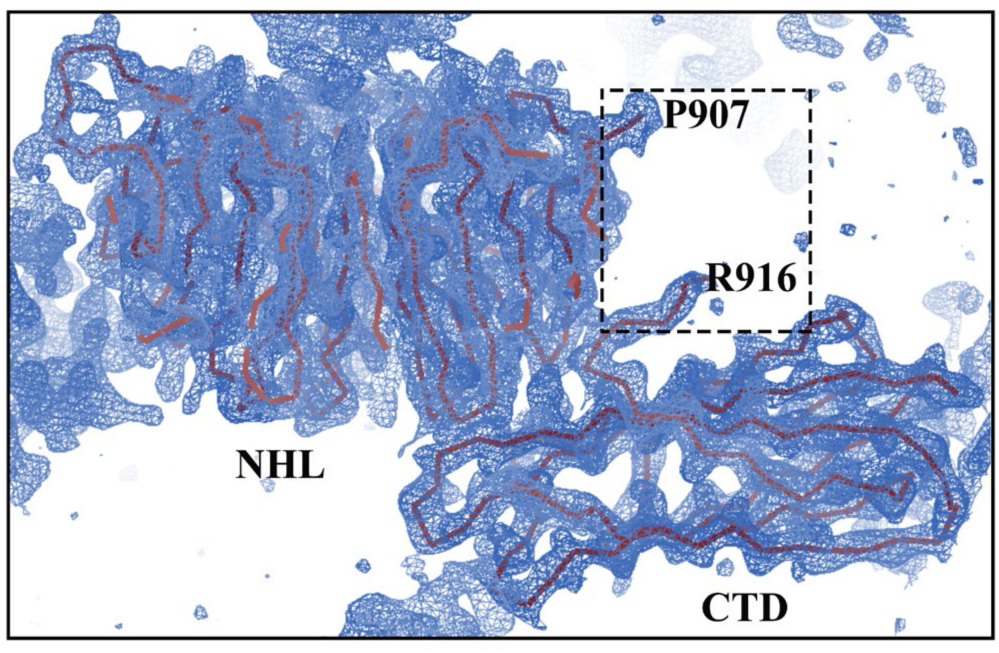
Electron density diagram of SOQ1_NHL(E859K)-CTD._ The 2Fo-Fc electron density map contoured at 1.0 σ are shown as blue mesh. The P907 and R916 are indicated, between which the density of eight residues are missing.

**Supplementary Figure S9.**
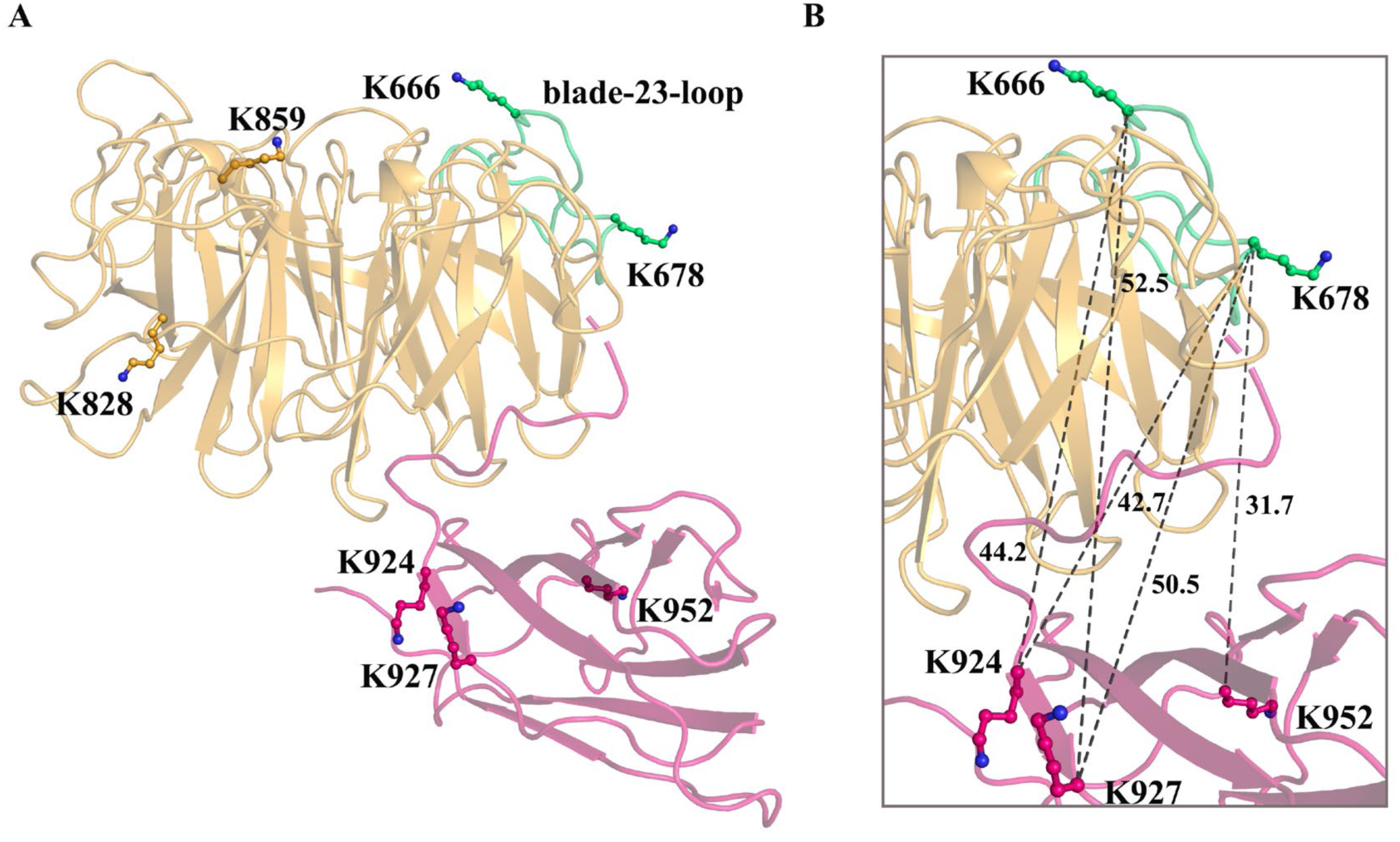
The location of cross-linked Lys residues in SOQ1_NHL(E859K)-CTD_ structure. (A) The overall structure of SOQ1_NHL(E859K)-CTD_, with the NHL domain and CTD colored bright orange and hot pink, respectively. The cross-linked residues are shown in stick-and-ball mode and labeled. The blade-23-loop is shown in green. (B) The cross-linked Lys pairs are linked with black dashed lines, and the distances (Å) between two Lys residues are indicated.

**Supplementary Figure S10.**
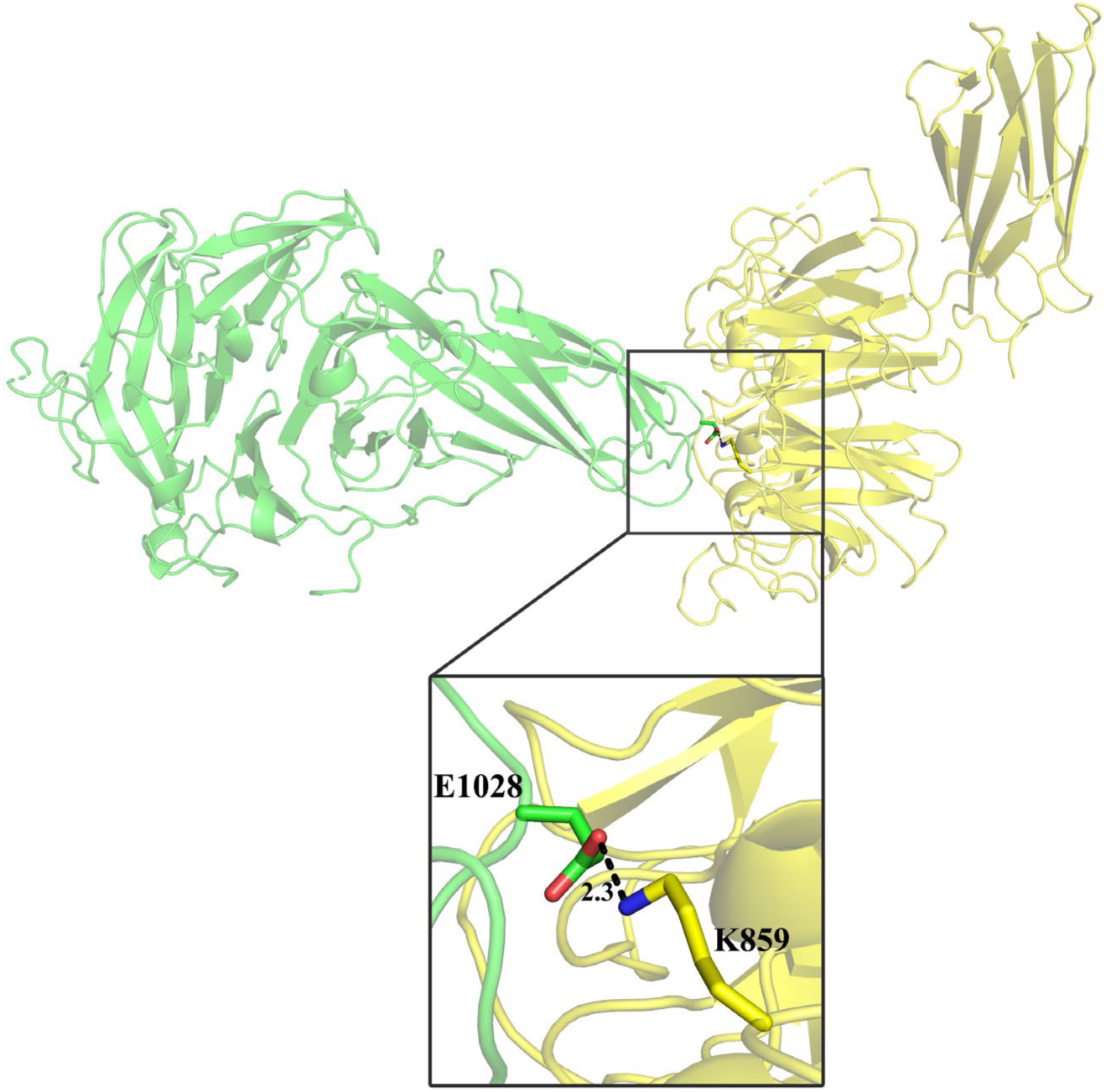
Molecular packing of SOQ1_NHL(E859K)-CTD_ in the crystal. Two symmetrically related SOQ1_NHL(E859K)-CTD_ molecules are shown in yellow and green, respectively. The residue K859 from one molecule strongly interacts with E1028 from another molecule, thus contributing to the crystal packing.

**Supplementary Figure S11.**
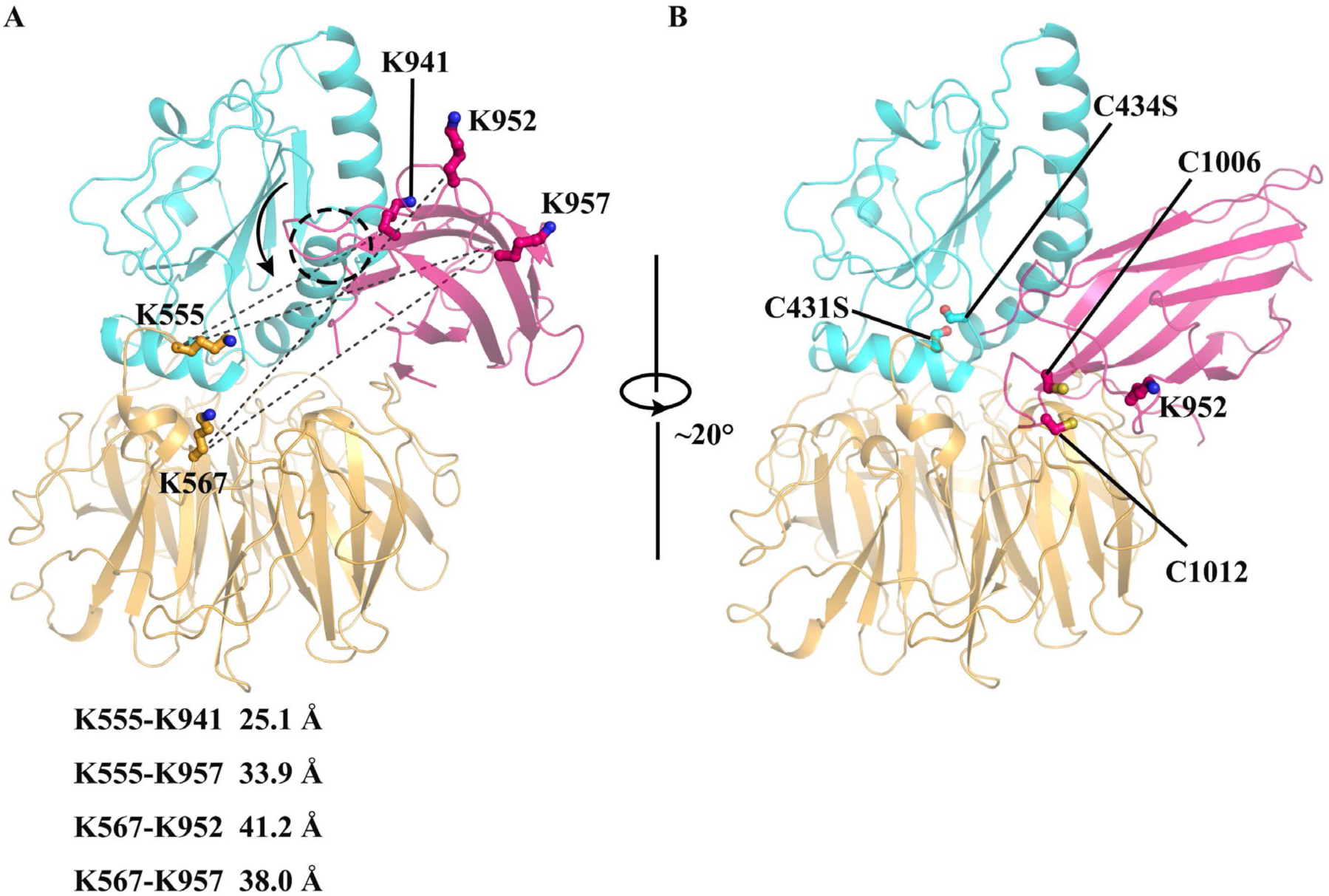
Structural model of SOQ1 lumenal domains. (A) Structural model of SOQ1 lumenal domains by superposing SOQ1_Trx(mut)-NHL_ and SOQ1_NHL-CTD_ structures, aligned on their NHL domain. The Trx-like domain, NHL domain and CTD are colored in cyan, bright orange and hot pink, respectively. The black dashed circle highlights the region where α2 of Trx-like domain overlaps with βB and βI of CTD. The residues K941, K952 and K957 from CTD are cross-linked with residues K555 and K567 from the TN-loop. The cross-linked pairs are linked by black dashed lines in the superposed structural model with distances shown below. The potential movement of CTD is indicated by arrow. (B) The modified model of SOQ-LD based on the cross-linking results. The C431S, C434S, C1006, C1012 and K952 are shown in stick-and-ball mode and labeled. Note that we used CTD from SOQ1_NHL(E859K)-CTD_ structure for modeling because it contains the complete βG-H loop with fully built residues C1006 and C1012.

**Supplementary Figure S12.**
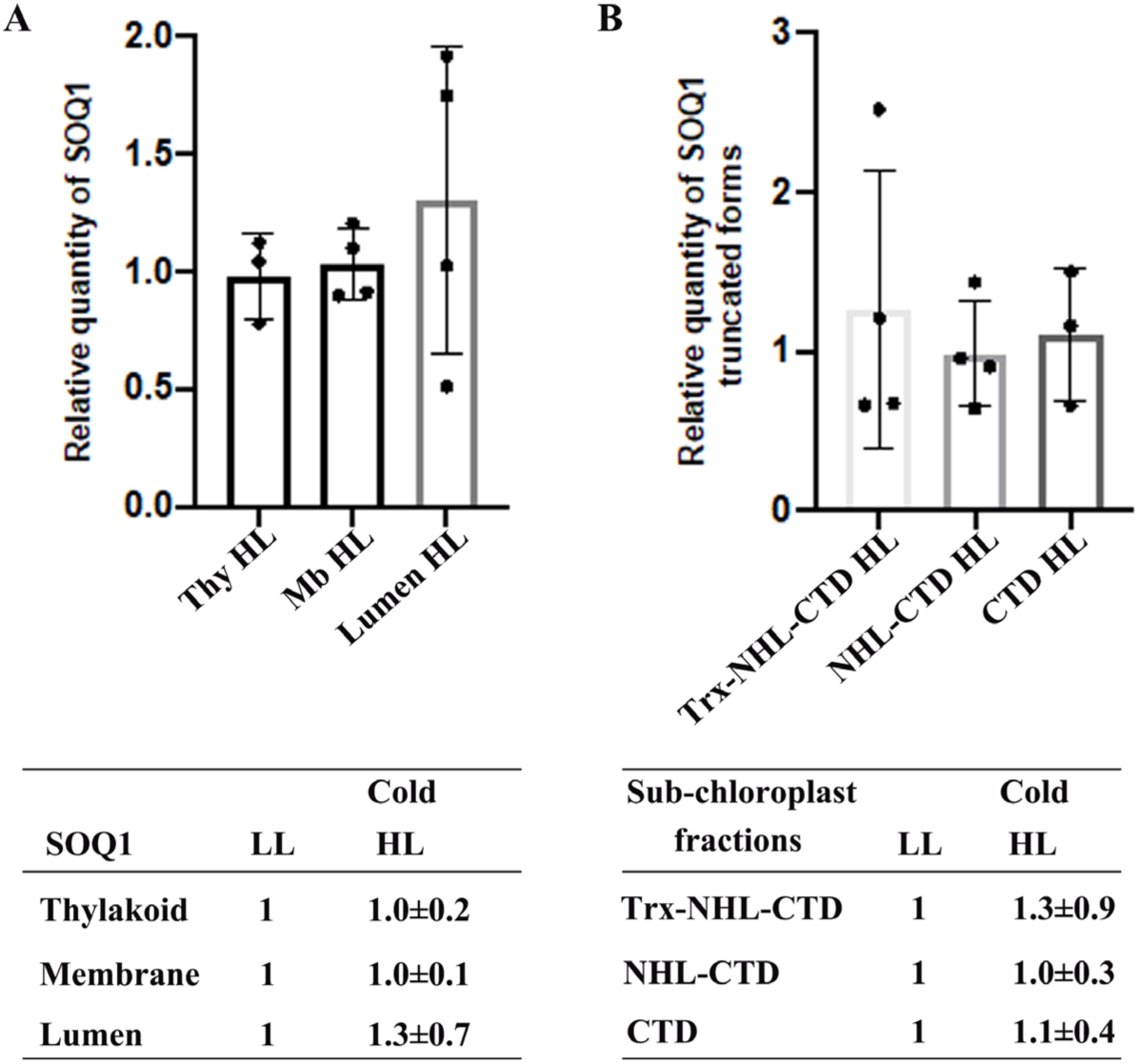
The accumulation of SOQ1 and its truncated forms is similar in stress and non-stress conditions. The relative quantities of SOQ1 in different fractionated thylakoid membrane preparations (**A**) and its truncated forms in the lumen (**B**). The quantities of SOQ1 in cold and high light (HL) were identified as a relative content to low light (LL). Thylakoid (Thyl), membrane (Mb), and lumen samples (full-length and truncated forms) isolated from Col-0 were loaded at the same amount of total protein. n = 4 independent biological experiments (n = 3 for CTD quantification).

**Table.S1.**
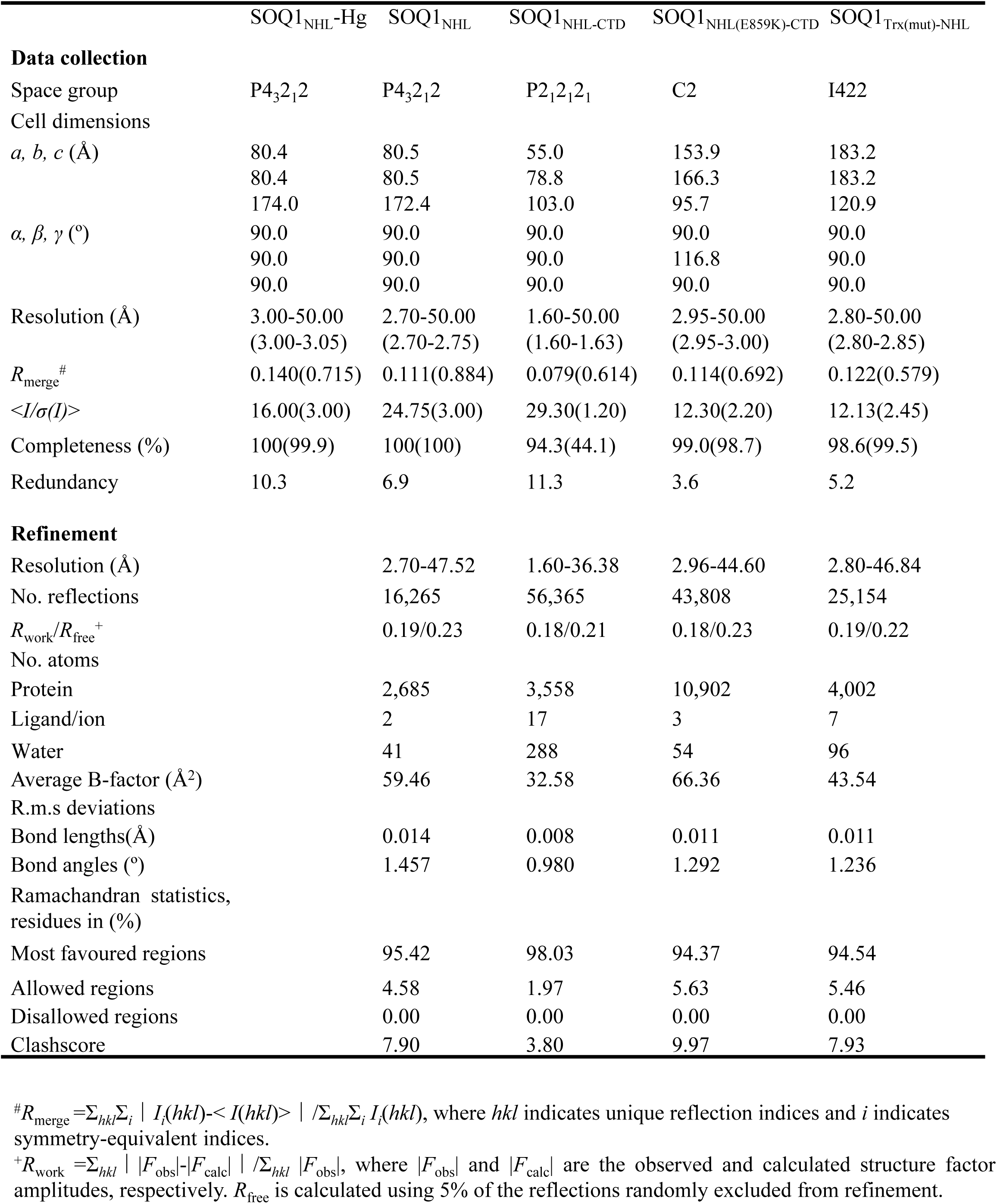
Data collection and refinement statistics.

|  | SOQ1<sub>NHL</sub>-Hg | SOQ1<sub>NHL</sub> | SOQ1<sub>NHL</sub>-CTD | SOQ1<sub>NHL(E859K)</sub>-CTD | SOQ1<sub>Trx(mut)</sub>-NHL |
| --- | --- | --- | --- | --- | --- |
| **Data collection** |  |  |  |  |  |
| Space group | P4<sub>3</sub>2<sub>1</sub>2 | P4<sub>3</sub>2<sub>1</sub>2 | P2<sub>1</sub>2<sub>1</sub>2<sub>1</sub> | C2 | I422 |
| Cell dimensions |  |  |  |  |  |
| *a*, *b*, *c* (Å) | 80.4 | 80.5 | 55.0 | 153.9 | 183.2 |
|  | 80.4 | 80.5 | 78.8 | 166.3 | 183.2 |
|  | 174.0 | 172.4 | 103.0 | 95.7 | 120.9 |
| *α*, *β*, *γ* (°) | 90.0 | 90.0 | 90.0 | 90.0 | 90.0 |
|  | 90.0 | 90.0 | 90.0 | 116.8 | 90.0 |
|  | 90.0 | 90.0 | 90.0 | 90.0 | 90.0 |
| Resolution (Å) | 3.00-50.00 | 2.70-50.00 | 1.60-50.00 | 2.95-50.00 | 2.80-50.00 |
|  | (3.00-3.05) | (2.70-2.75) | (1.60-1.63) | (2.95-3.00) | (2.80-2.85) |
| *R*<sub>merge</sub><sup>#</sup> | 0.140(0.715) | 0.111(0.884) | 0.079(0.614) | 0.114(0.692) | 0.122(0.579) |
| <*I*/*σ*(*I*)> | 16.00(3.00) | 24.75(3.00) | 29.30(1.20) | 12.30(2.20) | 12.13(2.45) |
| Completeness (%) | 100(99.9) | 100(100) | 94.3(44.1) | 99.0(98.7) | 98.6(99.5) |
| Redundancy | 10.3 | 6.9 | 11.3 | 3.6 | 5.2 |
| **Refinement** |  |  |  |  |  |
| Resolution (Å) |  | 2.70-47.52 | 1.60-36.38 | 2.96-44.60 | 2.80-46.84 |
| No. reflections |  | 16,265 | 56,365 | 43,808 | 25,154 |
| *R*<sub>work</sub>/*R*<sub>free</sub><sup>+</sup> |  | 0.19/0.23 | 0.18/0.21 | 0.18/0.23 | 0.19/0.22 |
| No. atoms |  |  |  |  |  |
| Protein |  | 2,685 | 3,558 | 10,902 | 4,002 |
| Ligand/ion |  | 2 | 17 | 3 | 7 |
| Water |  | 41 | 288 | 54 | 96 |
| Average B-factor (Å<sup>2</sup>) |  | 59.46 | 32.58 | 66.36 | 43.54 |
| R.m.s deviations |  |  |  |  |  |
| Bond lengths(Å) |  | 0.014 | 0.008 | 0.011 | 0.011 |
| Bond angles (°) |  | 1.457 | 0.980 | 1.292 | 1.236 |
| Ramachandran statistics,<br>residues in (%) |  |  |  |  |  |
| Most favoured regions |  | 95.42 | 98.03 | 94.37 | 94.54 |
| Allowed regions |  | 4.58 | 1.97 | 5.63 | 5.46 |
| Disallowed regions |  | 0.00 | 0.00 | 0.00 | 0.00 |
| Clashscore |  | 7.90 | 3.80 | 9.97 | 7.93 |

<sup>#</sup>*R*<sub>merge</sub> = $\sum_{hkl} \sum_i |I_i(hkl) - \langle I(hkl) \rangle| / \sum_{hkl} \sum_i I_i(hkl)$ , where *hkl* indicates unique reflection indices and *i* indicates symmetry-equivalent indices.
<sup>+</sup>*R*<sub>work</sub> = $\sum_{hkl} ||F_{\text{obs}}| - |F_{\text{calc}}|| / \sum_{hkl} |F_{\text{obs}}|$ , where *|F<sub>obs</sub>|* and *|F<sub>calc</sub>|* are the observed and calculated structure factor amplitudes, respectively. *R*<sub>free</sub> is calculated using 5% of the reflections randomly excluded from refinement.

**Table.S2.** BS3 cross-linking results of SOQ1-LD and SOQ1-LD(E859K).

| SOQ1-LD(K) |  | SOQ1-LD(E859K)(K) |  | Distance(Å) based on structure |  |
| --- | --- | --- | --- | --- | --- |
| NHL | CTD | NHL | CTD | SOQ1<sub>Trx(mut)-NHL</sub><br>+ SOQ1<sub>NHL-CTD</sub> | SOQ1<sub>Trx(mut)-NHL</sub><sup>+</sup><br>SOQ1<sub>NHL(E859K)-CTD</sub> |
| 666 | 924 | 666 | 924 | 19.1 | 43.8 |
| 666 | 927 | 666 | 927 | 9.5 | 52.3 |
| 678 | 924 | 678 | 924 | 22.3 | 42.4 |
| 678 | 927 | 678 | 927 | 15.6 | 50.3 |
| 678 | 941 |  |  | 28.9 |  |
| 678 | 952* | 678 | 952 | 41.9 | 31.4 |
|  |  | 675 | 924 |  | 41.3 |
|  |  | 828 | 924 |  | 38.1 |
|  |  | 844 | 952 |  | 52.3 |
|  |  | 859 | 924 |  | 37.0 |
|  |  | 859 | 927 |  | 46.2 |
| TN-Loop | CTD | TN-Loop | CTD |  |  |
| 567 | 927 | 567 | 927 | 23.8 | 38.2 |
| 567 | 952* | 567 | 952 | 41.2 | 39.5 |
| 567 | 957* | 567 | 957 | 38.0 | 33.9 |
| 555 | 941 |  |  | 25.1 |  |
| 555 | 957* |  |  | 33.9 |  |
|  |  | 567 | 924 |  | 30.1 |

\*: The distances of these cross-linking Lys-pairs are longer than the expected distance of approximately 24 Å, suggesting that CTD is tilted in solution and the region containing K952 and K957 is approaching to the Trx-NHL interface.

